# Tuning Hsp104 specificity to selectively detoxify α-synuclein

**DOI:** 10.1101/2020.04.15.043935

**Authors:** Korrie L. Mack, Hanna Kim, Meredith E. Jackrel, JiaBei Lin, Jamie E. DeNizio, Xiaohui Yan, Edward Chuang, Amber Tariq, Ryan R. Cupo, Laura M. Castellano, Kim A. Caldwell, Guy A. Caldwell, James Shorter

**Author notes:** Co-first author (Korrie L. Mack, Hanna Kim).

## Abstract

Hsp104 is an AAA+ protein disaggregase that solubilizes and reactivates proteins trapped in aggregated states. We have engineered potentiated Hsp104 variants to mitigate toxic misfolding of α-synuclein, TDP-43, and FUS implicated in fatal neurodegenerative disorders. Though potent disaggregases, these enhanced Hsp104 variants lack substrate specificity, and can have unfavorable off-target effects. Here, to lessen off-target effects, we engineer substrate-specific Hsp104 variants. By altering Hsp104 pore loops that engage substrate, we disambiguate Hsp104 variants that selectively suppress α-synuclein toxicity but not TDP-43 or FUS toxicity. Remarkably, α-synuclein-specific Hsp104 variants emerge that mitigate α-synuclein toxicity via distinct ATPase-dependent mechanisms, involving α-synuclein disaggregation or detoxification of α-synuclein conformers without disaggregation. Importantly, both types of α-synuclein-specific Hsp104 variant reduce dopaminergic neurodegeneration in a *C. elegans* model of Parkinson’s disease more effectively than non-specific variants. We suggest that increasing the substrate specificity of enhanced disaggregases could be applied broadly to tailor therapeutics for neurodegenerative disease.

## Introduction

There are no effective therapies for several fatal neurodegenerative diseases including Parkinson’s disease (PD), amyotrophic lateral sclerosis (ALS), and frontotemporal dementia (FTD). PD is marked by misfolding of α-synuclein (α-syn), a protein normally found in presynaptic terminals, which may function in synaptic vesicle recycling (Abeliovich and Gitler, 2016; Sun et al., 2019). α-Syn misfolds into toxic oligomers and amyloid fibrils, accumulating in characteristic Lewy bodies in the cytoplasm of dopaminergic neurons that degenerate in PD (Araki et al., 2019; Henderson et al., 2019; Shahmoradian et al., 2019). Indeed, protein misfolding and aggregation unite a spectrum of fatal neurodegenerative diseases (Chuang et al., 2018). Thus, it is important to develop therapeutics that directly antagonize the underlying toxic protein-misfolding events in neurodegenerative disease. In this way, proteins can be restored to their native conformation and function, which may halt the debilitating trajectory of neurodegeneration (March et al., 2019; Shorter, 2008, 2016, 2017).

Hsp104, an asymmetric, hexameric AAA+ protein disaggregase from yeast, is an intriguing therapeutic agent to directly target toxic protein-misfolding events in neurodegenerative disease (Jackrel and Shorter, 2015; Shorter and Southworth, 2019). Hsp104 uses energy from ATP binding and hydrolysis, and collaboration with Hsp70 and Hsp40, to reactivate misfolded proteins (DeSantis et al., 2012; Glover and Lindquist, 1998; Shorter and Southworth, 2019). Hsp104 hexamers are dynamic and adopt open “lock-washer” spiral states and closed ring structures that translocate polypeptides across the central channel (DeSantis et al., 2012; Gates et al., 2017; Michalska et al., 2019; Ye et al., 2019; Yokom et al., 2016). During protein disaggregation, pore-loop tyrosines grip substrate and ATP hydrolysis-driven conformational changes at the spiral seam ratchet substrate either partially or completely through the channel (Castellano et al., 2015; Gates et al., 2017; Lum et al., 2008; Lum et al., 2004; Sweeny et al., 2015; Tessarz et al., 2008; Ye et al., 2019, 2020). Thus, Hsp104 liberates individual polypeptides from soluble toxic oligomers, amorphous aggregates, and amyloid fibrils, which can then regain their functional form (DeSantis et al., 2012; Glover and Lindquist, 1998; Lo Bianco et al., 2008). Curiously, Hsp104 is not found in metazoa, but is conserved in eubacteria, algae, fungi, protozoa, and plants (Erives and Fassler, 2015). As humans lack an Hsp104 homolog, they have limited capacity to effectively counter overwhelming protein-misfolding events that underlie neurodegenerative disease (Shorter, 2011, 2017). Introduction of Hsp104 into animal models (e.g. worm, fly, mouse, and rat) protects against deleterious protein misfolding and neurodegeneration connected to PD and polyglutamine-expansion disorders (Cushman-Nick et al., 2013; Lo Bianco et al., 2008; Perrin et al., 2007; Satyal et al., 2000; Vacher et al., 2005). Nonetheless, the ability of Hsp104 to counter the misfolding and toxicity of human disease-linked proteins can be limited, requiring high Hsp104 concentrations (DeSantis et al., 2012; Lo Bianco et al., 2008).

To address this issue, we have engineered enhanced versions of Hsp104 that more effectively disaggregate disease-linked proteins such as α-syn (linked to PD), and TDP-43 and FUS (linked to ALS/FTD) under conditions where wild-type (WT) Hsp104 is ineffective (Jackrel et al., 2014a; Jackrel and Shorter, 2014a; Jackrel et al., 2015; Ryan et al., 2019; Tariq et al., 2019; Tariq et al., 2018; Torrente et al., 2016). Select potentiated variants reduce dopaminergic neurodegeneration in a *C. elegans* model of PD (Jackrel et al., 2014a) and reverse FUS aggregation and toxicity in mammalian cells (Yasuda et al., 2017). Typically, potentiated Hsp104 variants exhibit elevated ATPase activity, altered protomer cooperativity, altered substrate recognition, and prolonged substrate interactions, which enables more productive disaggregase activity (Durie et al., 2019; Jackrel et al., 2014a; Jackrel and Shorter, 2014a; Ye et al., 2020). Though powerful disaggregases, these potentiated Hsp104 variants can exhibit off-target toxicity, which may limit their progression along the therapeutic pipeline through more complex model systems (Jackrel et al., 2014a; Jackrel and Shorter, 2014b). This toxicity is likely caused by aberrant unfolding of essential substrates, resulting in unwanted off-target effects (Jackrel and Shorter, 2014a). Thus, enhanced substrate specificity is a desirable attribute for Hsp104 variants to effectively translate into higher organisms as therapeutics (Mack and Shorter, 2016).

Here, we hypothesized that specifically mutating Hsp104 residues known to contact substrate in a potentiated variant background would couple increased substrate-specificity to enhanced disaggregase activity. We found that specific alterations to pore-loop tyrosines that engage substrate directly endowed Hsp104 with the ability to selectively mitigate α-syn toxicity. Surprisingly, two classes of α-syn-specific Hsp104 variant emerged. The first class mitigated α-syn toxicity via ATPase-dependent disaggregation of α-syn inclusions. By contrast, an unanticipated second class mitigated α-syn toxicity via ATPase-dependent detoxification of α-syn conformers without disaggregation. Importantly, both types of α-syn-specific Hsp104 variant reduced dopaminergic neurodegeneration in a *C. elegans* model of PD more effectively than non-specific Hsp104 variants. Thus, we establish a new concept: specializing protein disaggregases against individual disease-associated substrates can improve their therapeutic utility. We anticipate that this concept can be applied broadly to diverse protein disaggregases and specific neurodegenerative disease proteins. In this way, specific toxic misfolding events could be remediated with tailor-made therapeutic disaggregases.

## Results

### Targeting Hsp104 pore-loop tyrosines through a rational engineering approach

Hsp104 consists of an N-terminal domain (NTD), nucleotide-binding domain 1 (NBD1), a middle domain (MD), nucleotide-binding domain 2 (NBD2), and a C-terminal domain (CTD) (Figure 1A) (Sweeny and Shorter, 2016). Hsp104 forms an asymmetric, ring-like hexamer, and threads substrate through its central channel, which is lined with substrate-binding pore loops (Figure 1B-D) (Gates et al., 2017; Lum et al., 2008; Tessarz et al., 2008; Yokom et al., 2016). Each NBD contains a highly-conserved tyrosine embedded within a pore-loop motif: ‘KYKG’ in NBD1 (residues 256-259) comprises pore-loop 1, and ‘GYVG’ in NBD2 (residues 661-664) comprises pore-loop 2 (Figure 1A-D) (Gates et al., 2017; Lum et al., 2008; Tessarz et al., 2008; Yokom et al., 2016). The pore-loop tyrosines, Y257 in pore-loop 1 and Y662 in pore-loop 2, are essential for disaggregation and substrate threading (DeSantis et al., 2012; Gates et al., 2017; Lum et al., 2008; Lum et al., 2004; Tessarz et al., 2008; Yokom et al., 2016). The pore loops are arranged in the Hsp104 axial channel as a ‘spiral staircase,’ allowing for substrate to be successfully translocated through the channel (Figure 1B-D) (Gates et al., 2017; Yokom et al., 2016). Y257 and Y662 contact substrate directly (Gates et al., 2017). We reasoned that subtly changing the properties of these highly-conserved tyrosines would enable us to tune the substrate repertoire of Hsp104. We altered each pore-loop tyrosine to a series of hydrophobic, aromatic, or uncharged polar residues to preserve different features of the original tyrosine residue, which was found to be optimal for protein disaggregation (Jackrel et al., 2014a; Lum et al., 2004). Using this rational engineering approach, we isolated α-syn-specific Hsp104 variants.

**Figure 1.**
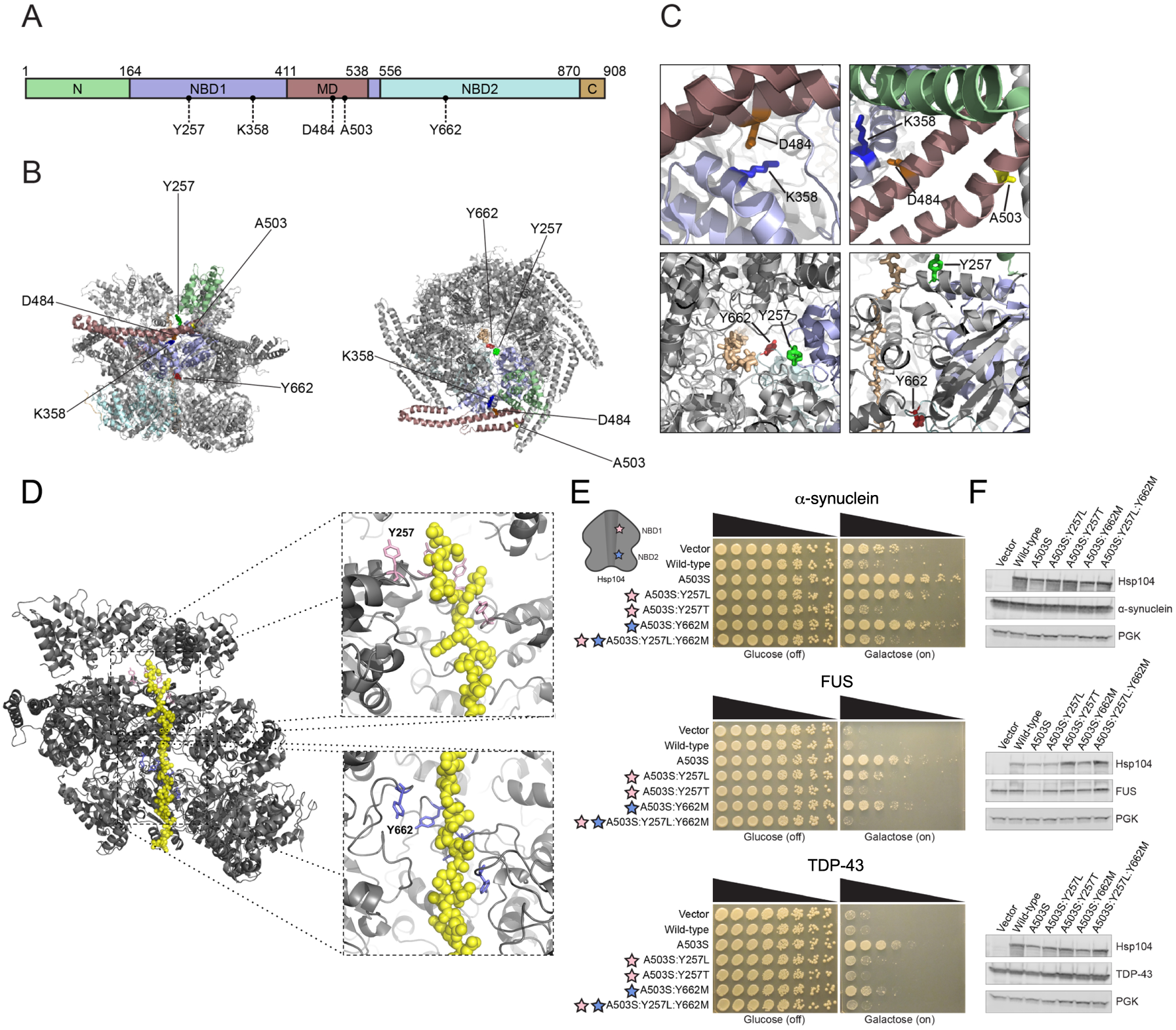
Pore-loop substitutions in Hsp104^A503S^ do not generate substrate-specific Hsp104 variants. **(A)** Domain map of Hsp104 showing N-terminal domain (N; green), nucleotide-binding domain 1 (NBD1; purple), middle domain (MD; maroon), nucleotide-binding domain 2 (NBD2; light blue), and C-terminal domain (C, orange). Location of residues assessed in this work are shown. **(B)** Structure of Hsp104 hexamer (side-view; left, top-down view; right) with residues from this study indicated. Colors of domains in protomer correspond to (A). PDB: 5VY9. **(C)** Zooms into Hsp104 structure highlighting K358:D484 interaction, and location of these residues in relation to A503. Substrate-binding pore-loop tyrosine residues are shown (Y257 and Y662) relative to polypeptide (casein) substrate (tan) in the Hsp104 channel. PDB: 5VY9. **(D)** Structure of Hsp104 hexamer (gray) with two subunits omitted to reveal polypeptide (casein) substrate (yellow) in the Hsp104 channel. Pore loops (pink; Y257 in NBD1 and purple; Y662 in NBD2) line the channel of Hsp104 in a “spiral staircase” manner. Pore-loop residues are essential for substrate translocation, as they establish the main contacts between Hsp104 and substrate. PDB: 5VY9. **(E)** Δ*hsp104* yeast integrated with α-syn-YFP (top), FUS (middle), or TDP-43 (bottom) on a galactose-inducible promoter were transformed with Hsp104 variants or an empty vector control. Yeast were spotted onto glucose (uninducing, off) and galactose (inducing, on) media in a five-fold serial dilution. Stars indicate substitution to pore loop in NBD1 (pink), NBD2 (purple). **(F)** Integrated strains from (E) were induced in the presence of Hsp104 variants or empty vector control for 5 hours (FUS, TDP-43) or 8 hours (α-syn). Yeast were lysed and processed for Western blot. 3-Phosphoglycerate kinase (PGK) is a loading control. See also **Figure S1, S2**, and **S3**.

### Mutating pore-loop tyrosines in Hsp104^A503S^ does not yield α-syn-specific variants

Mutating pore-loop tyrosines in WT Hsp104 does not enable suppression of α-syn, TDP-43, or FUS toxicity in yeast (Jackrel et al., 2014a). Thus, we first set out to engineer substrate-specific Hsp104 variants using a generally potentiated Hsp104 variant, Hsp104^A503S^ (Jackrel et al., 2014a), as a starting scaffold. Our goal was to leverage the elevated activity of Hsp104^A503S^ as a starting point to introduce substitutions that alter Hsp104 substrate selectivity. We assessed Hsp104 variant activity in powerful yeast models of α-syn, TDP-43, and FUS proteinopathy, which faithfully recapitulate several aspects of neurodegenerative disease, including protein aggregation and toxicity (Johnson et al., 2008; Outeiro and Lindquist, 2003; Sun et al., 2011). Importantly, these valuable yeast models have enabled identification of genetic suppressors and drug candidates that mitigate neurodegeneration in *C. elegans*, fly, mouse, rat, and human patient-derived neuronal models of disease (Becker et al., 2017; Chung et al., 2013; Cooper et al., 2006; Elden et al., 2010; Jackrel et al., 2014a; Jackson et al., 2015; Khurana et al., 2017; Tardiff et al., 2013).

Hsp104^A503S^ potently suppressed α-syn, TDP-43, and FUS toxicity in yeast (Figure 1E) (Jackrel et al., 2014a). In an effort to confer substrate specificity, we first introduced hydrophobic mutations to pore-loop 1 Y257 or pore-loop 2 Y662 to maintain the hydrophobic character of the tyrosine. Thus, we substituted a leucine at pore-loop 1 Y257. Although Y257 is highly conserved, leucine is also found rarely at this position in ∼0.5% of Hsp104 homologues according to our Generative Regularized ModeLs of proteINs (GREMLIN) analysis of 5,812 Hsp104 species variants (Ovchinnikov et al., 2014). Relative to Hsp104^A503S^, Hsp104^A503S:Y257L^ displayed reduced activity, as it suppressed α-syn and FUS toxicity, but not TDP-43 toxicity in yeast (Figure 1E). Though substitution of a hydrophobic residue at pore-loop 1 Y257 reduced toxicity suppression, introducing a hydrophobic methionine residue at pore-loop 2 Y662 had a different effect. Hsp104^A503S:Y662M^ suppressed toxicity of α-syn, TDP-43, and FUS, but not as strongly as Hsp104^A503S^ (Figure 1E). Combining these pore-loop mutations in Hsp104^A503S:Y257L:Y662M^ diminished any toxicity-suppression activity (Figure 1E).

Since hydrophobic substitutions at each pore loop did not lead to enhanced substrate specificity, we evaluated the effect of an uncharged, polar variant at Y257, and so introduced a threonine substitution. Interestingly, Hsp104^A503S:Y257T^ was unable to suppress toxicity of α-syn, TDP-43, or FUS (Figure 1E). Expressing each pore-loop variant did not noticeably affect expression levels of disease-associated substrates (Figure 1F). Altogether, slightly altering Hsp104 pore-loop tyrosines in the potentiated Hsp104^A503S^ background did not generate any variants with the desired substrate specificity. Indeed, several pore-loop mutations diminished the ability of Hsp104^A503S^ to mitigate toxicity.

### Hsp104^K358D^ can be tailored via tuning pore loops for substrate specificity

Hsp104^A503S^ is potentiated via disruption of interprotomer contacts between helix L1 and helix L3 of the MD (Gates et al., 2017; Heuck et al., 2016; Tariq et al., 2019; Ye et al., 2020), but was not amenable for tuning substrate specificity (Figure 1E). Thus, we wondered whether substrate selectivity could be more effectively engineered in a Hsp104 variant potentiated by a different mechanism than Hsp104^A503S^. We first considered Hsp104 with a K358D mutation in NBD1 as a starting scaffold. Hsp104^K358D^ has a mutation that disrupts interactions between D484 in the MD and K358 in NBD1 (Figure 1C) similar to the previously-studied Hsp104^K358E^ (Gates et al., 2017; Lipinska et al., 2013). Hsp104^K358E^ has elevated ATPase activity relative to Hsp104, but has been reported to be extremely toxic to yeast (Lipinska et al., 2013). By contrast, Hsp104^K358D^ was not toxic to Δ*hsp104* yeast at 30°C (Figure S1). Thus, we explored whether this Hsp104 variant could suppress toxicity of neurodegenerative disease-linked substrates, and whether we could now tune pore-loop residues to engender substrate specificity.

Hsp104^K358D^ suppressed α-syn, FUS, and TDP-43 toxicity in yeast (Figure 2A). Thus, Hsp104^K358D^ can antagonize proteotoxicity connected to PD and ALS/FTD. Using Hsp104^K358D^ as a starting scaffold, we systematically altered the substrate-binding, pore-loop tyrosines, Y257 and Y662, as with Hsp104^A503S^. Interestingly, Hsp104^K358D:Y257L^ suppressed toxicity of α-syn, FUS, and TDP-43 more effectively than Hsp104^K358D^ (Figure 2A). Thus, a leucine at position 257 in pore-loop 1 is more productive than a tyrosine in the Hsp104^K358D^ background in striking contrast to Hsp104^A503S^ (Figure 1E, 2A). Interestingly, Hsp104^K358D:Y662L^, with a leucine at pore-loop 2 Y662, did not suppress FUS or TDP-43 toxicity, but mildly suppressed α-syn toxicity (Figure 2A). Thus, increased α-syn selectivity can originate by altering Y662 in the K358D background. We next explored hydrophobic, aromatic substitutions at pore loops 1 and 2. Hsp104^K358D:Y257F/W^ and Hsp104^K358D:Y662F/W^ suppressed toxicity of α-syn, FUS, and TDP-43 (Figure 2A). We also substituted hydrophobic residues isoleucine and valine at the pore-loop tyrosines. Hsp104^K358D:Y257I/V^ slightly suppressed α-syn toxicity, and did not suppress FUS or TDP-43 toxicity (Figure 2A). Thus, α-syn selectivity can also emerge by mutating Y257 in the K358D background. By contrast, Hsp104^K358D:Y662I/V^ did not suppress α-syn, FUS, or TDP-43 toxicity (Figure 2A). Interestingly, even a minor change from leucine to isoleucine at the 662 position resulted in diminished ability to suppress α-syn toxicity.

**Figure 2.**
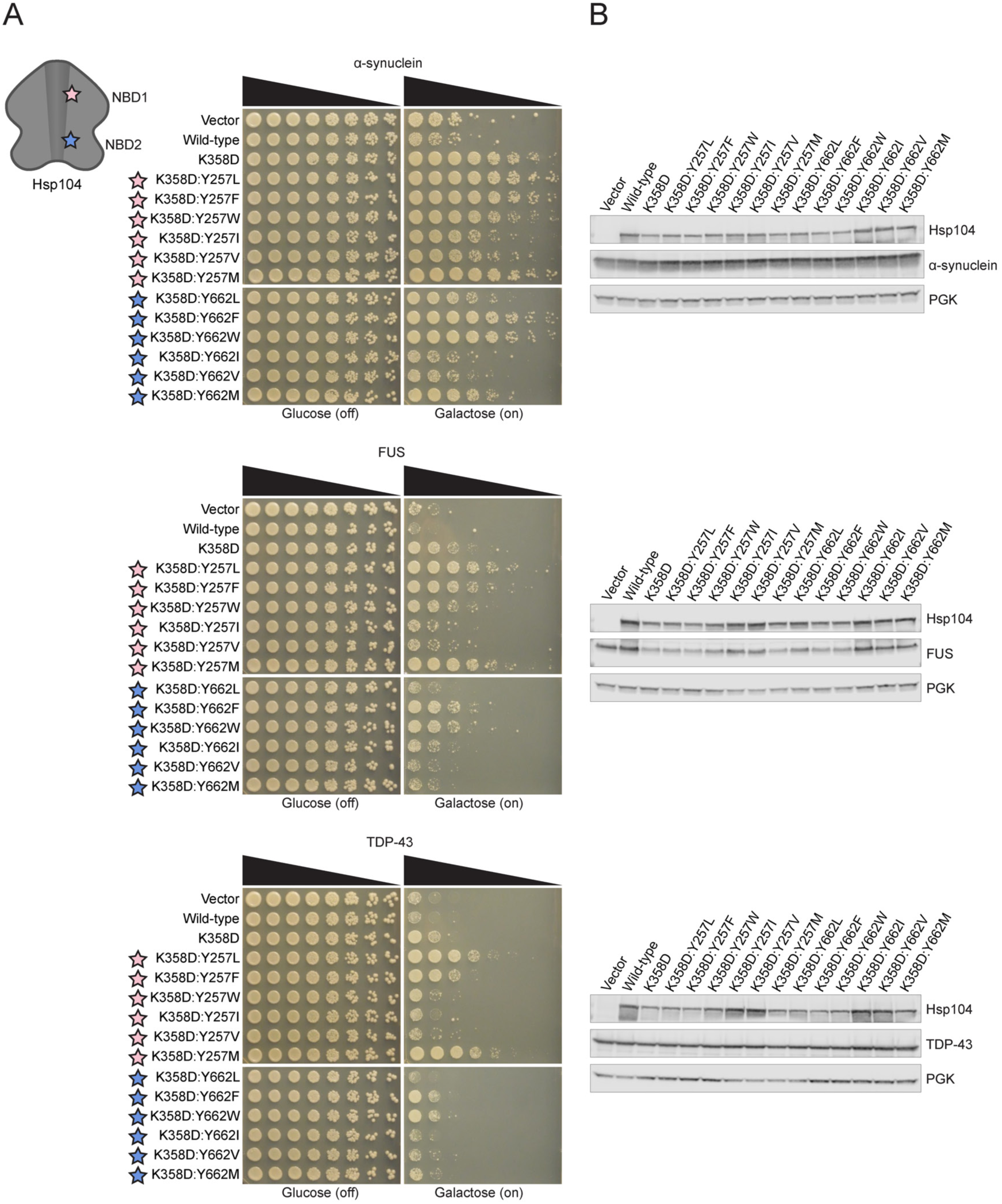
Hsp104^K358D:Y662M^ selectively suppresses α-syn toxicity. **(A)** Δ*hsp104* yeast integrated with α-syn-YFP (top), FUS (middle), or TDP-43 (bottom) on a galactose-inducible promoter were transformed with Hsp104 variants or an empty vector control. Yeast were spotted onto glucose (uninducing, off) and galactose (inducing, on) media in a five-fold serial dilution. Stars indicate substitution to pore loop in NBD1 (pink), NBD2 (purple). **(B)** Integrated strains from (A) were induced in the presence of Hsp104 variants or empty vector control for 5 hours (FUS, TDP-43) or 8 hours (α-syn). Yeast were lysed and lysates visualized via Western blot. 3-Phosphoglycerate kinase (PGK) is a loading control. See also **Figure S1, S2**, and **S3**.

Generally, our pore-loop Hsp104 variants did not grossly affect α-syn or TDP-43 expression levels (Figure 2B). A subset of these variants mildly reduced FUS expression (Figure 2B), but we have shown before that this mild reduction in FUS levels is not required to suppress FUS toxicity by enhanced Hsp104 variants (Jackrel and Shorter, 2014a; Tariq et al., 2019). Collectively, our data suggest that engineered pore-loop variants do not suppress disease-associated substrate toxicity by severely lowering substrate levels.

Given the changes in toxicity suppression when a single pore-loop tyrosine was mutated in the Hsp104^K358D^ background, we next tested the effect of double pore-loop tyrosine mutations. As Hsp104^K358D:Y257L^ showed robust suppression of α-syn, FUS, and TDP-43 toxicity, we used this variant as a starting scaffold to tune the second pore-loop tyrosine. Hsp104^K358D:Y257L:Y662L^ suppressed toxicity of α-syn, but only slightly reduced FUS and TDP-43 toxicity (Figure 3A). Hsp104^K358D:Y257L:Y662F/W^ suppressed toxicity of all three substrates, consistent with the finding that phenylalanine and tryptophan substitutions at either pore loop alone do not affect the activity of enhanced Hsp104 variants (Jackrel et al., 2014a) (Figure 3A). Interestingly, introducing polar, uncharged residues (S, T, Q, N) at pore-loop 2 Y662 ablated the toxicity suppression activity of Hsp104^K358D:Y257L^, again suggesting that pore-loop 2 Y662 is not as amenable to mutation as pore-loop 1 Y257 (Figure 3A). Overall, the double pore-loop Hsp104 variants were similar to those with single pore-loop mutations in that aside from a set of variants that mildly reduced FUS expression, the variants did not grossly affect disease-associated substrate levels (Figure 3B).

**Figure 3.**
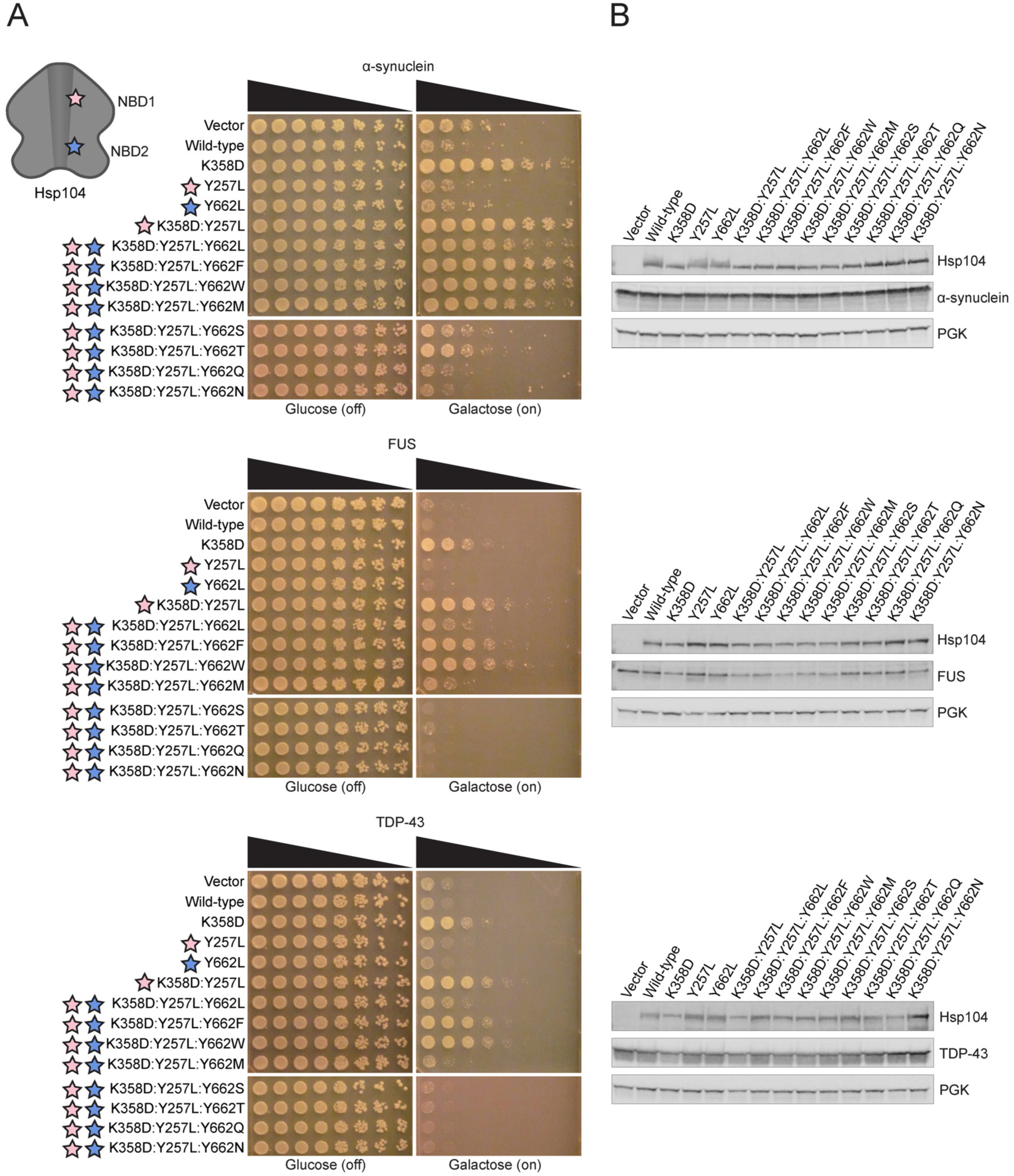
Hsp104^K358D:Y257L:Y662M^ selectively suppresses α-syn toxicity. **(A)** Δ*hsp104* yeast integrated with α-syn-YFP (top), FUS (middle), or TDP-43 (bottom) on a galactose-inducible promoter were transformed with Hsp104 variants or an empty vector control. Yeast were spotted onto glucose (uninducing, off) and galactose (inducing, on) media in a five-fold serial dilution. Stars indicate substitution to pore loop in NBD1 (pink), NBD2 (purple). **(B)** Integrated strains from (A) were induced in the presence of Hsp104 variants or empty vector control for 5 hours (FUS, TDP-43) or 8 hours (α-syn). Yeast were lysed and lysates visualized via Western blot. 3-Phosphoglycerate kinase (PGK) is a loading control. See also **Figure S3**.

Our rational approach to altering the substrate repertoire of Hsp104 revealed several trends for pore-loop substitutions that are favorable for general potentiation. In the enhanced Hsp104^K358D^ background, aromatic residues (W, F) at Y257 or Y662 maintained enhanced toxicity suppression activity relative to WT Hsp104 (Figure 2A). Substituting aromatic residues for Y662 in the potentiated Hsp104^K358D:Y257L^ background also maintained potentiation relative to WT Hsp104 (Figure 3A). Hydrophobic residues (I, V) at Y257 in the Hsp104^K358D^ background were not favorable for potentiated activity against TDP-43 or FUS, but did yield more selective variants that mildly suppressed α-syn toxicity (Figure 2A). Additionally, uncharged, polar residues at Y662 in Hsp104^K358D:Y257L^ eliminated potentiation (Figure 3A).

### Y662M in pore-loop 2 of Hsp104^K358D^ or Hsp104^K358D:Y257L^ confers α-syn selectivity

We unearthed further α-syn-specific variants by substituting a Met residue at the NBD2 pore-loop tyrosine. Thus, a Y662M substitution at pore-loop 2 in the Hsp104^K358D^ or Hsp104^K358D:Y257L^ created variants that selectively suppressed α-syn toxicity (Figure 2A, 3A). Indeed, Hsp104^K358D:Y662M^ and Hsp104^K358D:Y257L:Y662M^ suppressed α-syn toxicity, but were ineffective against FUS or TDP-43 toxicity (Figure 2A, 3A). These variants did not grossly affect α-syn expression levels (Figure 2B, 3B). Interestingly, introducing a Met residue at pore-loop 1 (Y257) in the Hsp104^K358D^ background did not confer substrate specificity, but rather strengthened suppression of α-syn, FUS, and TDP-43 toxicity (Figure 2A). Thus, replacing Tyr with Met in pore-loop 1 versus pore-loop 2 has distinct effects on the substrate selectivity of Hsp104.

### Hsp104^D484K^ can also be tailored via tuning pore loops for substrate specificity

In addition to Hsp104^K358D^, Hsp104^D484K^ is another Hsp104 variant that disrupts MD-NBD1 interactions normally mediated by K358 in NBD1 and D484 in helix L2 of the MD (Figure 1C) (Lipinska et al., 2013). To our surprise, Hsp104^D484K^ was not toxic to yeast at 30°C (Figure S1). Thus, neither Hsp104^K358D^ nor Hsp104^D484K^ are overtly toxic to yeast at 30°C. We generated a set of pore-loop Hsp104^D484K^ variants, which displayed very similar effects on α-syn, FUS, and TDP-43 toxicity and expression to those studied in the Hsp104^K358D^ background (Figure S2A, B). For example, like Hsp104^K358D:Y257L/M^, Hsp104^D484K:Y257L/M^ enhanced activity against α-syn, FUS, and TDP-43 (Figure S2A). Furthermore, mutation of Y257 to I or V, or Y662 to L or M yielded Hsp104^D484K^ variants that selectively suppressed α-syn toxicity (Figure S2A). Thus, in potentiated backgrounds created by disrupting NBD1 and MD contacts (e.g. K358D and D484K) we establish general rules for: (a) mutating pore loops to enhance suppression of α-syn, FUS, and TDP-43 toxicity; and (b) mutating pore loops to selectively suppress α-syn toxicity. Specifically, enhanced activity is conferred by Y257L/M mutations and α-syn-selectivity is conferred by Y257I/V mutation or Y662L/M mutations.

### Y257T/Q in pore-loop 1 of Hsp104^K358D^ confer α-syn selectivity

We next introduced alternative uncharged, polar residues Thr, Gln, Ser, or Asn at pore-loop 1, Y257. As these substitutions still maintain the polar, uncharged character of tyrosine we reasoned they might tune the substrate repertoire of Hsp104. Remarkably, Hsp104^K358D:Y257T/Q^ specifically suppressed α-syn toxicity and did not suppress FUS or TDP-43 toxicity (Figure 4A). Furthermore, Hsp104^K358D:Y257T/Q^ did not reduce α-syn expression levels (Figure 4B). Interestingly, Hsp104^K358D:Y257S/N^ did not effectively suppress α-syn, TDP-43, or FUS toxicity (Figure 4A). We focused on Hsp104^K358D:Y257T^ as a representative among the uncharged, polar (Hsp104^K358D:Y257T/Q^) and hydrophobic (Hsp104^K358D:Y257I/V^) α-syn-specific variants. This variant suppressed α-syn toxicity to a lesser extent than Hsp104^K358D:Y662M^ (Figure 2A, 4A), revealing that even among α-syn-specific variants, we can tune the degree of toxicity suppression.

**Figure 4.**
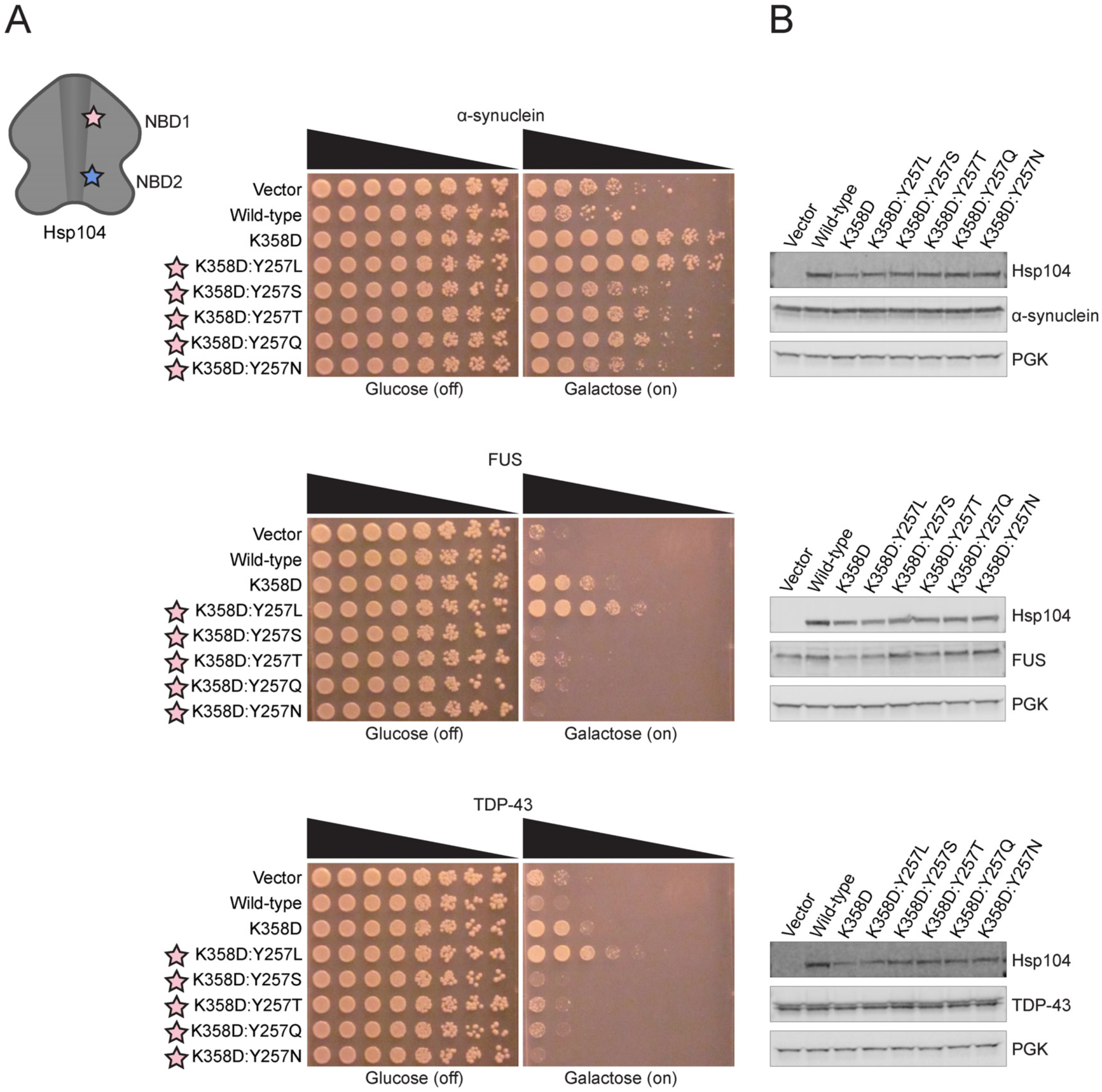
Hsp104^K358D:Y257T/Q^ selectively suppress α-syn toxicity. **(A)** Δ*hsp104* yeast integrated with α-syn-YFP (top), FUS (middle), or TDP-43 (bottom) on a galactose-inducible promoter were transformed with Hsp104 variants or an empty vector control. Yeast were spotted onto glucose (uninducing, off) and galactose (inducing, on) media in a five-fold serial dilution. Stars indicate substitution to pore loop in NBD1 (pink), NBD2 (purple). **(B)** Integrated strains from (A) were induced in the presence of Hsp104 variants or empty vector control for 5 hours (FUS, TDP-43) or 8 hours (α-syn). Yeast were lysed and lysates visualized via Western blot. 3-Phosphoglycerate kinase (PGK) is a loading control. See also **Figure S3**.

### α-Syn-specific Hsp104 variants suppress α-syn toxicity through an ATPase-dependent mechanism

We next determined whether α-syn-specific Hsp104 variants suppressed α-syn toxicity via an ATP hydrolysis-dependent process as for other potentiated Hsp104 variants (Jackrel et al., 2014a; Torrente et al., 2016). Thus, we made double Walker A (DWA) mutations (K218T and K620T) that render Hsp104 unable to bind ATP at NBD1 and NBD2 (DeSantis et al., 2012; Franzmann et al., 2011; Parsell et al., 1991; Schaupp et al., 2007; Schirmer et al., 1998; Torrente et al., 2016). The DWA mutations inhibited potentiated variants Hsp104^A503S^, Hsp104^K358D^, and Hsp104^K358D:Y257L^ from suppressing toxicity of α-syn, FUS, and TDP-43 (Figure S3A). DWA mutations in Hsp104^K358D:Y257T^ diminished suppression of α-syn toxicity (Figure S3A), suggesting Hsp104^K358D:Y257T^ also employs an ATP hydrolysis-dependent mechanism. Hsp104^K358D:Y662M^ and Hsp104^K358D:Y257L:Y662M^ were also inactivated by DWA mutations (Figure S3A). Thus, these α-syn-specific Hsp104 variants also work through a mechanism reliant on ATP hydrolysis. The reduction in therapeutic Hsp104 activity conferred by DWA mutations was not due to reduced Hsp104 levels (Figure S3B).

### α-Syn-specific Hsp104 variants do not suppress FUS aggregation

We next evaluated the ability of the α-syn-specific Hsp104 variants to antagonize FUS aggregation in yeast. α-Syn-specific Hsp104 variants should not affect FUS aggregates in yeast, as FUS is no longer recognized as a substrate. In ALS/FTD and in yeast proteinopathy models, FUS mislocalizes to cytoplasmic aggregates (Couthouis et al., 2011; Ju et al., 2011; Sun et al., 2011). Enhanced Hsp104 variants, Hsp104^K358D^ and Hsp104^K358D:Y257L^, effectively suppressed FUS aggregation compared to Hsp104, which had no effect (Figure S4A, B). In agreement with our toxicity-suppression findings, α-syn-specific variants Hsp104^K358D:Y257T^, Hsp104^K358D:Y662M^, and Hsp104^K358D:Y257L:Y662M^ did not antagonize FUS aggregation (Figure S4A, B). Thus, our α-syn-specific variants do not suppress FUS toxicity or aggregation in yeast, strongly advocating for tailored substrate specificity.

### α-Syn-specific Hsp104 variants do not restore TDP-43 to the nucleus

As with FUS, we expect α-syn-specific Hsp104 variants should not suppress mislocalization of TDP-43, as these variants have an altered substrate repertoire. TDP-43 is normally localized to the nucleus in human cells, but mislocalizes to cytoplasmic aggregates in ALS/FTD pathology, which is recapitulated in yeast (Elden et al., 2010; Figley and Gitler, 2013; Johnson et al., 2008; Johnson et al., 2009). While Hsp104 was unable to return TDP-43 to the nucleus, enhanced variants, Hsp104^K358D^ and Hsp104^K358D:Y257L^, significantly restored nuclear localization to TDP-43, as ∼39% and ∼50% of cells respectively harbored nuclear TDP-43 (Figure S4C, D). Importantly, α-syn-specific variants Hsp104^K358D:Y662M^ and Hsp104^K358D:Y257L:Y662M^ did not restore TDP-43 to the nucleus, and α-syn-specific variant Hsp104^K358D:Y257T^ had only a slight effect on TDP-43 localization (Figure S4C, D). Thus, α-syn-specific Hsp104 variants do not suppress TDP-43 toxicity or restore TDP-43 to the nucleus, consistent with increased substrate selectivity.

### α-Syn-specific Hsp104 variants differentially affect cytoplasmic α-syn foci

α-Syn initially localizes to the plasma membrane in yeast, but eventually forms toxic cytoplasmic inclusions that are detergent-insoluble, contain high molecular weight α-syn species, react with the amyloid-diagnostic dye Thioflavin-S, and cluster cytoplasmic vesicles reminiscent of aspects of Lewy pathology in PD (Araki et al., 2019; Gitler et al., 2008; Jackrel et al., 2014a; Outeiro and Lindquist, 2003; Shahmoradian et al., 2019; Soper et al., 2008; Tenreiro et al., 2014; Zabrocki et al., 2005). While Hsp104 is unable to antagonize formation of cytoplasmic α-syn foci and return α-syn to the plasma membrane, enhanced variants Hsp104^K358D^ and Hsp104^K358D:Y257L^ robustly clear α-syn foci (Figure 5A, B). Approximately 62% of yeast cells expressing Hsp104^K358D^ have plasma membrane-localized α-syn and ∼48% of cells expressing Hsp104^K358D:Y257L^ have plasma membrane-localized α-syn, compared to only ∼18% cells when Hsp104 is present (Figure 5B). α-Syn-specific variants Hsp104^K358D:Y662M^ and Hsp104^K358D:Y257L:Y662M^ also eradicated α-syn foci (Figure 5A, B). Unexpectedly, α-syn-specific variant Hsp104^K358D:Y257T^ did not significantly eliminate α-syn foci relative to Hsp104 (Figure 5A, B). Thus, Hsp104^K358D:Y257T^ is likely not suppressing α-syn toxicity by resolving α-syn foci. The uncoupling of toxicity suppression and clearance of α-syn foci is a novel feature of Hsp104^K358D:Y257T^, as we have not previously engineered Hsp104 in this way. Thus, Hsp104^K358D:Y257T^ emerges as a new, substrate-specific Hsp104 variant that is distinct from other α-syn-specific variants Hsp104^K358D:Y662M^ and Hsp104^K358D:Y257L:Y662M^.

**Figure 5.**
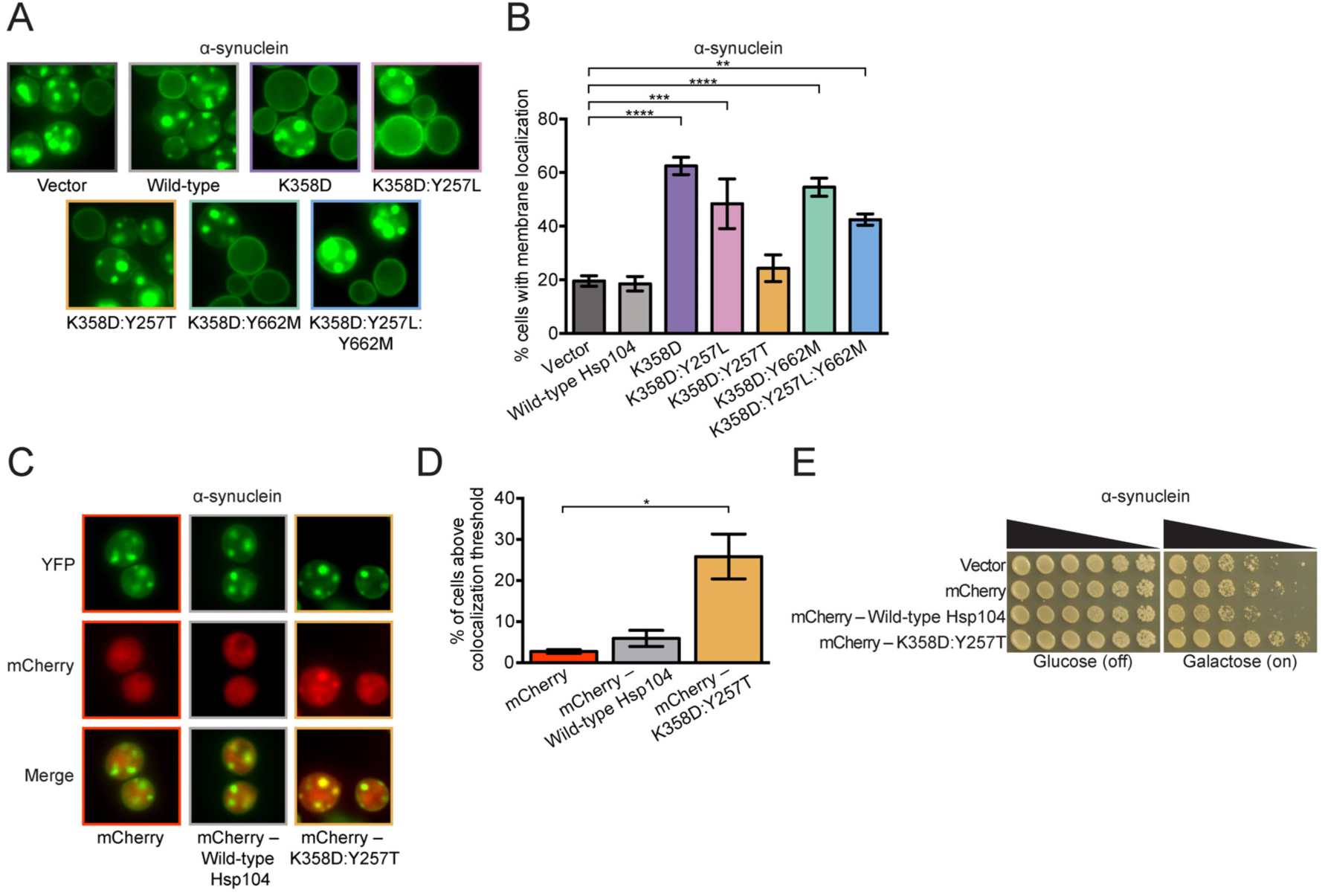
α-Syn-specific Hsp104 variants differentially eradicate α-syn inclusions. **(A)** Representative fluorescence microscopy images of Δ*hsp104* yeast integrated with α-syn-YFP and transformed with Hsp104 variants or an empty vector control. **(B)** Localization of α-syn-YFP in yeast was quantified by counting the number of cells with plasma membrane-localized α-syn-YFP or α-syn-YFP foci in the cytoplasm. Values represent means ± SEM (n = 3-6). \*\**P*<0.01, \*\*\**P*<0.001, \*\*\*\**P*<0.0001; One-way ANOVA with Dunnett’s *post-hoc* test. **(C)** Representative fluorescence microscopy images of Δ*hsp104* yeast integrated with α-syn-YFP and transformed with mCherry – Hsp104 variants or mCherry alone as a control. **(D)** Colocalization of mCherry– Hsp104 variants or mCherry alone with α-syn-YFP foci was quantified using a semi-automated procedure. Percentages of cells above the colocalization threshold are plotted. Values represent means ± SEM (n = 3-6). \**P*<0.05; One-way ANOVA with Dunnett’s *post-hoc* test. **(E)** Δ*hsp104* yeast integrated with α-syn-YFP on a galactose-inducible promoter were transformed with mCherry – Hsp104 variants or mCherry alone. Yeast were spotted onto glucose (uninducing, off) and galactose (inducing, on) media in a five-fold serial dilution. See also **Figure S4**.

### Hsp104^K358D:Y257T^ colocalizes with α-syn foci more frequently than Hsp104

As Hsp104^K358D:Y257T^ suppressed α-syn toxicity in yeast but did not eliminate α-syn foci, we were curious if this variant colocalized with α-syn foci. One explanation for the uncoupling of toxicity suppression and clearance of α-syn foci is that Hsp104^K358D:Y257T^ is still able to recognize and colocalize with α-syn foci to mitigate their toxicity. We tagged Hsp104 and Hsp104^K358D:Y257T^ with mCherry to visualize their cellular localization in yeast expressing α-syn-YFP. C-terminally mCherry-tagged Hsp104^K358D:Y257T^ more frequently colocalized with α-syn foci than mCherry-tagged Hsp104 or mCherry alone (Figure 5C, D). mCherry-Hsp104^K358D:Y257T^ colocalized with α-syn foci in ∼25% of cells (Figure 5D) and suppressed α-syn toxicity (Figure 5E). That mCherry-Hsp104^K358D:Y257T^ colocalizes with α-syn foci more than mCherry-Hsp104 suggests that Hsp104^K358D:Y257T^ may remodel α-syn conformers to a less toxic conformation or otherwise quench their toxicity (e.g. by shielding surfaces from interacting with other essential cell components, such as intracellular vesicles). Furthermore, Hsp104^K358D:Y257T^ likely engages α-syn foci more efficiently than Hsp104, as it has enhanced substrate specificity.

### α-Syn-specific Hsp104 variants have altered ATPase activity

To elucidate the underlying mechanism for substrate-specific toxicity suppression, we surveyed various biochemical properties of a representative set of Hsp104 variants. We first tested the ATPase activity to determine if α-syn-specific variants differ in ability to hydrolyze ATP relative to Hsp104 and potentiated variants. We purified the generally potentiated MD variant, Hsp104^A503S^, and the generally potentiated NBD1 variant, Hsp104^K358D^, both of which exhibit elevated ATPase activity compared to Hsp104 (Figure 6A) (Jackrel et al., 2014a; Lipinska et al., 2013; Ye et al., 2020). We also purified a generally potentiated pore-loop variant, Hsp104^K358D:Y257L^, and three α-syn specific variants, Hsp104^K358D:Y257T^, Hsp104^K358D:Y662M^, and Hsp104^K358D:Y257L:Y662M^. Interestingly, Hsp104^K358D:Y257L^ exhibited reduced ATPase activity compared to Hsp104^K358D^, but slightly higher ATPase activity than Hsp104 (Figure 6A). Thus, Hsp104^K358D:Y257L^ provides another example of a potentiated Hsp104 that does not have greatly elevated basal ATPase activity as with Hsp104^D504C^ (Jackrel et al., 2014a). Nonetheless, this effect was unexpected as mutating Y257 in WT Hsp104 does not typically reduce basal ATPase activity (DeSantis et al., 2012; Lum et al., 2008; Torrente et al., 2016). This reduction in ATPase activity was even more pronounced for the α-syn-specific variant Hsp104^K358D:Y257T^, which had ATPase activity similar to Hsp104, but reduced ATPase activity compared to Hsp104^K358D^ (Figure 6A). Thus, the K358D mutation sensitizes Hsp104 ATPase activity to Y257 mutations, perhaps due to destabilization of the small domain of NBD1 similar to another enhanced NBD1 variant, Hsp104^E360R^, which also bears a charge-reversing mutation in the vicinity of K358 (Tariq et al., 2019; Ye et al., 2020).

**Figure 6.**
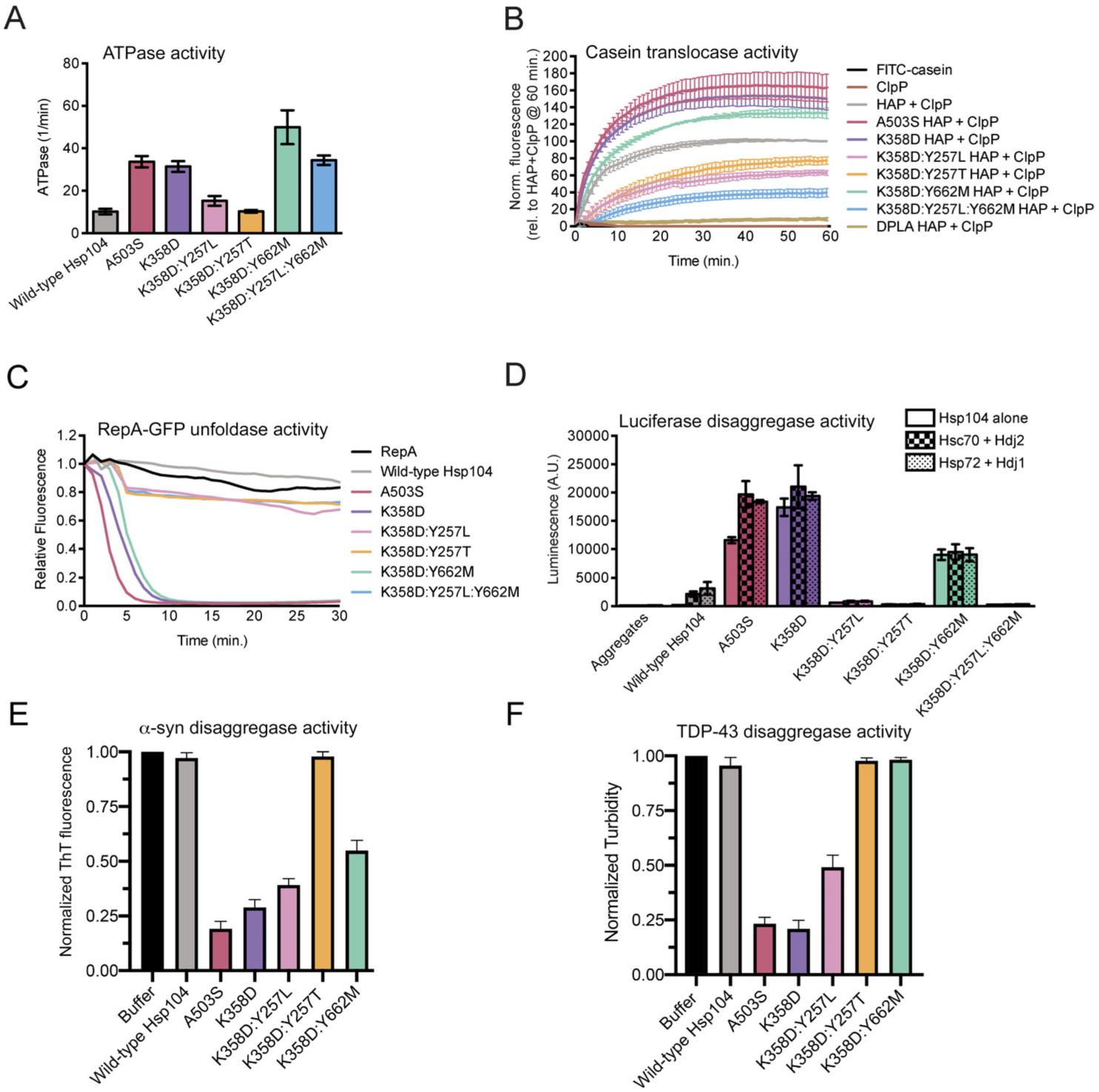
α-Syn-specific Hsp104 variants have distinct biochemical properties. **(A)** ATPase activity of Hsp104 variants. Values represent means ± SEM (n=4). **(B)** HAP variants (0.167µM hexamer) plus ClpP (21µM monomer) were incubated with FITC-casein, and FITC-casein degradation was measured by fluorescence. Negative controls FITC-casein alone (black) and ClpP alone control (gray) included. Values were normalized to HAP plus ClpP at 60 minutes and represent means ± SEM (n = 3-6). **(C)** RepA_1-25_-GFP (0.7µM) was incubated with Hsp104 variants (2.1µM hexamer) plus GroEL^trap^ (2.5µM tetradecamer), and RepA_1-25_-GFP unfolding was measured by fluorescence. Negative control of RepA_1-25_-GFP alone included. A representative trial is shown (n = 3). **(D)** Hsp104 variants alone (1µM hexamer, solid bars) or with chaperone pairs Hsc70 and Hdj2 (0.167µM each, checkered bars), or Hsp72 and Hdj1 (0.167µM each, dotted bars), were incubated with urea-denatured luciferase aggregates. Reactivation activity was measured by luminescence. Negative control of luciferase aggregates alone included. Values represent means ± SEM (n = 2-8). **(E)** α-Syn fibrils (3μM monomer) were incubated without or with Hsp104, Hsp104^A503S^, Hsp104^K358D^, Hsp104^K358D:Y257T^ or Hsp104^K358D:Y662M^ (3μM) plus Hsc70 (3μM), Hdj1 (3μM), ATP (20 mM) and an ATP regeneration system (20mM creatine phosphate and 0.5μM creatine kinase) for 2h at 30°C. Disaggregation was assessed by Thioflavin-T (ThT) fluorescence. Values represent means ± SEM (n = 3). **(F)** TDP-43 fibrils (3 μM monomer) were incubated without or with Hsp104, Hsp104^A503S^, Hsp104^K358D^, Hsp104^K358D:Y257T^ or Hsp104^K358D:Y662M^ (3μM) plus Hsc70 (3μM), Hdj1 (3μM), ATP (20 mM) and an ATP regeneration system (20mM creatine phosphate and 0.5μM creatine kinase) for 2h at 30°C. Disaggregation was assessed by turbidity (absorbance at 350nm). Values represent means ± SEM (n = 3). See also **Figure S5**.

The other α-syn-specific variants, Hsp104^K358D:Y662M^ and Hsp104^K358D:Y257L:Y662M^, exhibited ATPase activity equal to or greater than the activity of Hsp104^A503S^ and Hsp104^K358D^ (Figure 6A). Indeed, the Y662M mutation stimulated ATPase activity in the K358D background, and even counteracted the effect of the Y257L mutation such that Hsp104^K358D:Y257L:Y662M^ had higher ATPase activity than Hsp104^K358D:Y257L^ and similar ATPase activity to Hsp104^K358D^ (Figure 6A). This result was also unanticipated as Y662 mutations do not typically affect WT Hsp104 ATPase activity (DeSantis et al., 2012; Lum et al., 2004; Torrente et al., 2016), indicating that the K358D mutation sensitizes Hsp104 ATPase activity to Y662 mutations.

The difference in ATPase activity between α-syn-specific variants is a key biochemical distinction between them. These findings further suggest that α-syn-specific variants likely employ different toxicity-suppression mechanisms. Thus, the lower ATPase activity of Hsp104^K358D:Y257T^ may limit clearance of α-syn foci in yeast, whereas elevated ATPase activity of Hsp104^K358D:Y662M^ and Hsp104^K358D:Y257L:Y662M^ likely enables clearance of α-syn foci.

### α-Syn-specific Hsp104 variants have altered translocase activity

Next, we assessed translocase activity of each Hsp104 variant. Hsp104 translocates substrates across its central channel during disaggregation (DeSantis et al., 2012; Shorter and Southworth, 2019). To assess substrate translocation, we evaluated each Hsp104 variant in the HAP background *in vitro*. HAP is an Hsp104 variant with three missense mutations (G739I:S740G:K741F) that enable interaction with the chambered protease, ClpP (Tessarz et al., 2008). Thus, HAP but not Hsp104 can translocate substrate into the proteolytic chamber of ClpP where it is degraded. We monitored translocation of a model unfolded substrate fluorescein isothiocyanate (FITC)-labeled casein through HAP via degradation by ClpP, which liberates FITC and increases fluorescence (DeSantis et al., 2012). Mutating each pore-loop Tyr to Ala (Y257A:Y662A), which ablates substrate translocation and disaggregation (DeSantis et al., 2012; Lum et al., 2008; Tessarz et al., 2008; Torrente et al., 2016), resulted in the DPLA (‘double pore-loop alanine’) HAP variant that lacks translocation activity (Figure 6B). Thus, minimal passive translocation occurs in this system. We previously found that HAP^A503V^ translocated substrate more rapidly than HAP (Jackrel et al., 2014a). Likewise, HAP^A503S^ and HAP^K358D^ translocated substrate more rapidly than HAP (Figure 6B). This accelerated substrate translocation may promote enhanced disaggregase activity.

Interestingly, α-syn-specific variants exhibited distinct activities. HAP^K358D:Y662M^ translocated FITC-casein more effectively than HAP, whereas HAP^K358D:Y257L:Y662M^ was less effective (Figure 6B). In the context of Hsp104, these variants have similar ATPase activity (Figure 6A). Thus, HAP^K358D:Y257L:Y662M^ likely has reduced ability to grip FITC-casein resulting in decelerated translocation. HAP^K358D:Y257L^ and HAP^K358D:Y257T^ translocated FITC-casein at a similar rate, but were less effective than HAP (Figure 6B). These findings suggest that interfering with Y257 of pore-loop 1 likely weakens grip of casein and thereby reduces translocase activity. Changes in substrate grip and translocase activity likely contribute to altered patterns of substrate specificity, as they could enable fine-tuning of the force used by Hsp104 to disaggregate different substrates. Indeed, Hsp104 variants with low translocase activity may only partially translocate substrate, thus changing the way a substrate is disaggregated.

### α-Syn-specific Hsp104 variants have altered unfoldase activity

We next evaluated the unfoldase activity of each Hsp104 variant using RepA_1-25_-GFP as a model substrate. In these reactions, we included GroEL^trap^, which captures newly unfolded RepA_1-25_-GFP and prevents its refolding (Weber-Ban et al., 1999). Hsp104 is unable to unfold RepA_1-25_-GFP under these conditions, whereas potentiated Hsp104 variants, Hsp104^A503S^ and Hsp104^K358D^, rapidly unfold RepA_1-25_-GFP (Figure 6C) (Jackrel et al., 2014a). Remarkably, α-syn-specific Hsp104 variants showed distinct unfoldase activity from one another. α-Syn-specific Hsp104^K358D:Y662M^ unfolded RepA_1-25_-GFP almost as rapidly as Hsp104^A503S^ and Hsp104^K358D^ (Figure 6C). By contrast, α-syn-specific Hsp104^K358D:Y257T^ and Hsp104^K358D:Y257L:Y662M^, as well as generally potentiated pore-loop variant Hsp104^K358D:Y257L^, only modestly unfolded RepA_1-25_-GFP (Figure 6C). Here too, it is likely that pore-loop 1 mutations weaken substrate gripping by Hsp104 and limit unfolding of the RepA_1-25_-GFP substrate. As Hsp104^K358D:Y662M^ possessed strong unfoldase activity similar to generally potentiated Hsp104 variants, enhanced substrate recognition or unfolding power may contribute to its mechanism of toxicity suppression. The stark difference in unfoldase activity between α-syn-specific Hsp104 variants is another distinction in their biochemical properties that could contribute to different modes of α-syn toxicity suppression in yeast.

### α-Syn-specific Hsp104 variants have altered luciferase disaggregase activity

Next, we assessed disaggregase activity of each Hsp104 variant against aggregated luciferase. Differences in the ability to reactivate this model substrate could help to illuminate general disaggregase properties of substrate-selective Hsp104 variants. Hsp104 was unable to reactivate disordered luciferase aggregates on its own, and required Hsp70 and Hsp40 (Figure 6D) (Glover and Lindquist, 1998; Jackrel et al., 2014a). Generally potentiated variants, Hsp104^A503S^ and Hsp104^K358D^, exhibited similar activities, and had high luciferase reactivation activity even in the absence of Hsp70 and Hsp40 (Figure 6D). Interestingly, the generally potentiated pore-loop variant Hsp104^K358D:Y257L^ was less active than Hsp104, indicating that Y257L mutation is inhibitory with respect to specifically disaggregating luciferase (Figure 6D). Indeed, it appears that Y257L alters activity against model substrates *in vitro*, but not against disease substrates in yeast. Of the three α-syn specific Hsp104 variants, only Hsp104^K358D:Y662M^ had luciferase reactivation activity (Figure 6D). Hsp104^K358D:Y662M^ was more active than Hsp104, and displayed comparable activity in the absence and presence of Hsp70 and Hsp40 (Figure 6D). Notably, each Hsp104 variant showed similar activity with two different human Hsp70 and Hsp40 pairs, Hsc70/Hdj2, and Hsp72/Hdj1, suggesting this activity is not dependent on the presence of specific Hsp70 or Hsp40 variants (Figure 6D). The distinct luciferase reactivation activity of Hsp104^K358D:Y662M^ and lack of activity of Hsp104^K358D:Y257L^, Hsp104^K358D:Y257T^, and Hsp104^K358D:Y257L:Y662M^, likely result from a specific effect on luciferase from perturbing pore-loop 1. Altering pore-loop 1 in the context of the enhanced Hsp104^K358D^ background must interfere with disaggregation of disordered luciferase aggregates. Indeed, these variants may no longer recognize luciferase effectively as a result of altered substrate-specificity.

### Hsp104^K358D:Y662M^ disaggregates α-syn fibrils but not TDP-43 fibrils

Next, we assessed the ability of Hsp104 variants to disaggregate preformed α-syn and TDP-43 fibrils (Jackrel et al., 2014a). Hsp104 displayed limited ability to dissociate α-syn or TDP-43 fibrils under these conditions, whereas Hsp104^A503S^ and Hsp104^K358D^ effectively dismantled α-syn and TDP-43 fibrils (Figure 6E, F). Hsp104^K358D:Y257L^ also effectively disaggregated α-syn and TDP-43 fibrils (Figure 6E, F). Thus, although ineffective against luciferase (Figure 6D), Hsp104^K358D:Y257L^ was active against α-syn and TDP-43 fibrils (Figure 6E, F). By contrast, Hsp104^K358D:Y257T^ was unable to dissolve α-syn or TDP-43 fibrils (Figure 6E, F). Thus, the ability of Hsp104^K358D:Y257T^ to reduce α-syn toxicity is separated from α-syn disaggregation. Finally, Hsp104^K358D:Y662M^ could disassemble α-syn fibrils, but not TDP-43 fibrils (Figure 6E, F), indicating a change in substrate specificity that explains the ability to reduce α-syn toxicity, but not TDP-43 toxicity.

### α-Syn-specific Hsp104 variants show distinct toxicity phenotypes at 37°C

As the α-syn-specific Hsp104 variants had differing biochemical profiles, we wondered if these variants showed differences in their off-target toxicity to yeast. Thus, we evaluated their toxicity at 37°C in the absence of neurodegenerative disease-associated substrate. Yeast are mildly stressed at 37°C as essential proteins are slightly unfolded. Potentiated Hsp104 variants can nonspecifically target key proteins that are slightly unfolded, resulting in a strong toxic phenotype when expressed in yeast (Jackrel and Shorter, 2014a). Thus, this assay can be used to uncover Hsp104 variants that exhibit less off-target toxicity that may be advanced to more complex model systems. Consistent with previous results (Jackrel et al., 2014a), the potentiated variants Hsp104^A503V^ and Hsp104^K358D^ were toxic to yeast relative to Hsp104 at 37°C but not 30°C (Figure S5A). Interestingly, generally potentiated Hsp104^K358D:Y257L^ and α-syn-specific Hsp104^K358D:Y257T^ virtually eliminated this toxic phenotype (Figure S5A). Low toxicity to yeast could result from reduced targeting of essential, unfolded substrates in yeast and enhanced recognition of toxic substrates. It appears that the Y257L mutation reduces Hsp104 activity against off-target yeast substrates, but not against disease substrates. The ability to suppress toxicity of neurodegenerative disease-associated substrates, coupled with very low inherent toxicity to yeast, are desirable attributes for translating these disaggregases into effective therapeutic agents. Surprisingly, however, both Hsp104^K358D:Y662M^ and Hsp104^K358D:Y257L:Y662M^ were noticeably toxic to yeast at 37°C (Figure S5A). We suggest these variants could be targeting a substrate under mild stress conditions that is essential for yeast. It seems the Y662M mutation reduces Hsp104 activity specifically against TDP-43 and FUS, but not against α-syn, or off-target yeast substrates.

### α-Syn-specific Hsp104 variants confer yeast thermotolerance to different extents

Next, we assessed the ability of α-syn-specific Hsp104 variants to confer thermotolerance in yeast. One of the central functions of Hsp104 is to help yeast survive heat shock by resolubilizing proteins that aggregate during stress (Lindquist and Kim, 1996; Parsell et al., 1994; Parsell et al., 1991; Sanchez and Lindquist, 1990; Sanchez and Taulien, 1992; Wallace et al., 2015). We hypothesized that substrate-specific Hsp104 variants should be defective in conferring thermotolerance, in that they are no longer able to recognize a broad array of substrates for solubilization. We evaluated each Hsp104 variant under the native heat shock element (HSE) Hsp104 promoter relative to Hsp104 in their ability to confer thermotolerance to yeast. We first induced Hsp104 expression at 37°C, then heat-shocked yeast at 50°C (Figure S5B). All Hsp104 variants were expressed at similar levels (Figure S5C). While an empty vector conferred no thermotolerance (very few yeast survive a 30 minute heat shock at 50°C), Hsp104 transformed into a Δ*hsp104* strain effectively complemented the yeast survival observed in yeast expressing Hsp104 endogenously (Figure S5B). Potentiated variants Hsp104^A503V^ and Hsp104^K358D^ did not retain full Hsp104 activity in conferring thermotolerance (Figure S5B). Likewise, Hsp104^K358D:Y662M^ and Hsp104^K358D:Y257L:Y662M^ also conferred reduced thermotolerance (Figure S5B). This reduced activity is perhaps due to off-target toxicity at the 37°C pretreatment step (Figure S5A). Interestingly, Hsp104^K358D:Y257L^ and Hsp104^K358D:Y257T^ conferred similar levels of reduced thermotolerance (Figure S5B). For these variants, which are not toxic at 37°C (Figure S5A), the reduced thermotolerance indicates that they do not disaggregate the complete repertoire of Hsp104 substrates (Figure S5B).

### Several α-syn-specific Hsp104 variants prevent α-syn-induced neurodegeneration in *C. elegans*

We next tested whether the Hsp104 variants that selectively suppressed α-syn toxicity in yeast can prevent neurodegeneration in a *C. elegans* model of PD, which has successfully validated PD-relevant modifiers of α-syn toxicity (Cao et al., 2005; Cooper et al., 2006; Harrington et al., 2010; Jackrel et al., 2014a; Tardiff et al., 2013). Preventing α-syn-induced neurodegeneration in this model is a key step in translating our findings in yeast to a full metazoan nervous system. We tested α-syn specific Hsp104^K358D:Y257T^, Hsp104^K358D:Y662M^, and Hsp104^K358D:Y257L:Y662M^, as well as generally potentiated Hsp104^K358D^ and Hsp104^K358D:Y257L^. In this transgenic model, expression of human α-syn and Hsp104 variants is driven selectively in the dopaminergic neurons of worms using the endogenous promoter of the dopamine transporter gene, P_*dat-1*_. When α-syn is expressed alone, ∼23% of worms retained the full complement of dopaminergic neurons at day 7, and ∼10% at day 10 post hatching (Figure 7A), which recapitulates α-syn neurotoxicity in human α-synucleinopathies. Hsp104 was unable to protect against α-syn-induced neurodegeneration (Figure 7A) (Jackrel et al., 2014a). Surprisingly, generally potentiated Hsp104^K358D^ and Hsp104^K358D:Y257L^ also did not significantly protect against neurodegeneration (Figure 7A). Moreover, the α-syn-specific Hsp104^K358D:Y257L:Y662M^ did not protect against neurodegeneration, suggesting that this variant also did not translate from yeast to worm.

**Figure 7.**
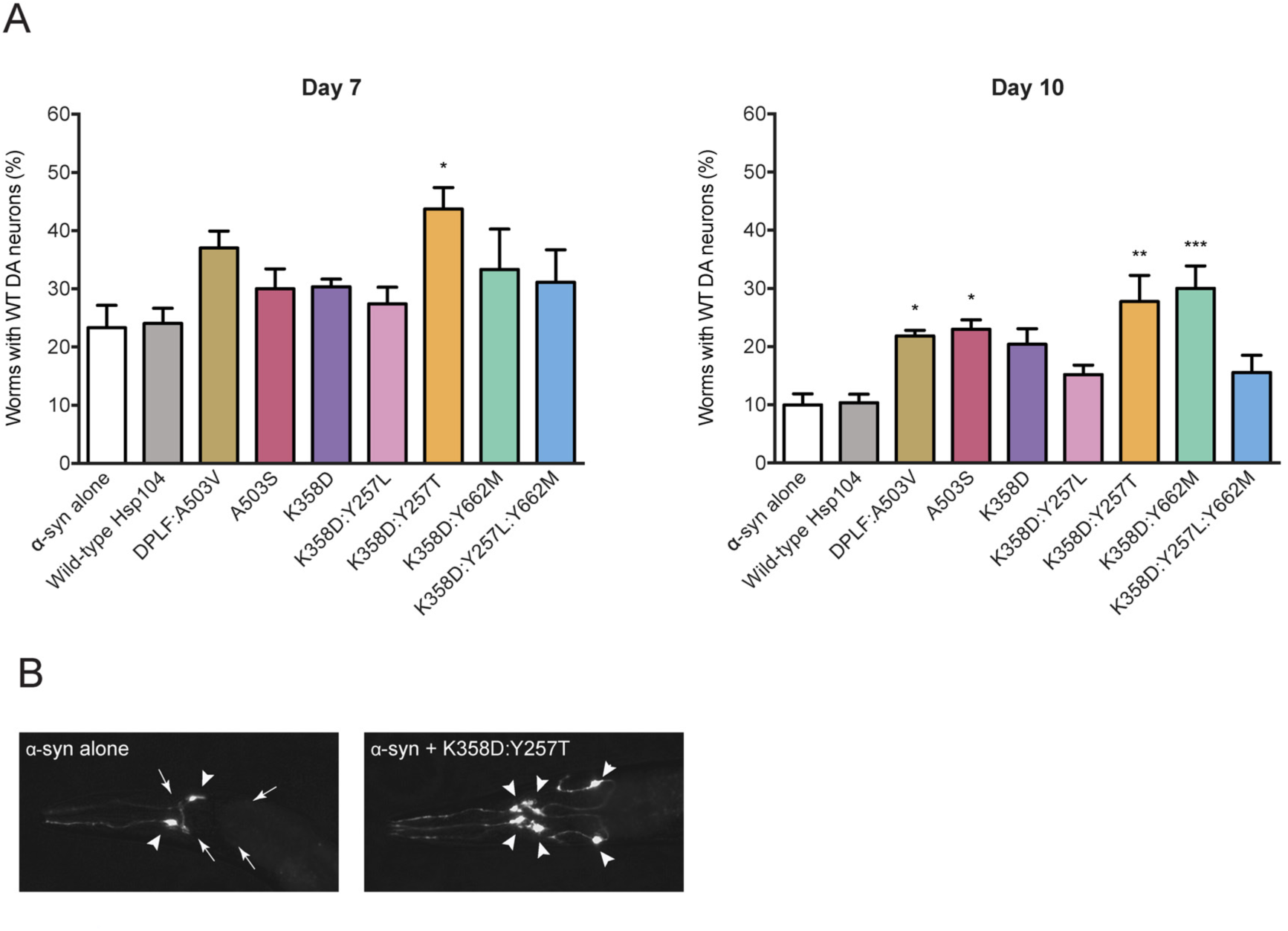
Several α-syn-specific Hsp104 variants prevent dopaminergic neuron degeneration in a *C. elegans* model of PD. **(A)** Hsp104 variants and α-syn were co-expressed in the dopaminergic (DA) neurons of *C. elegans*. Animals expressing Hsp104^K358D:Y257T^ are significantly protected against α-syn-induced DA neuron degeneration at day 7 post-hatching (left). DA neurodegeneration was exacerbated at day 10 post-hatching (right). Though not protective at day 7 post-hatching, overexpression of Hsp104 variants Hsp104^DPLF:A503V^, Hsp104^A503S^, and Hsp104^K358D:Y662M^ protected against α-syn-induced DA neurodegeneration at day 10 post-hatching (right). Notably, the Hsp104^K358D:Y257T^ variant consistently exhibits neuroprotection at both the earlier and later timepoints. Values represent mean ± SEM, n = 30 per replicate, three independent experiments were performed/variant and three distinct worm stable lines were generated for each Hsp104 variant. \**P*<0.05, \*\**P*<0.01, \*\*\**P*<0.001; One-way ANOVA with Dunnett’s *post-hoc* test. **(B)** A representative image of *C. elegans* DA neurons in worms expressing α-syn alone (left) or α-syn + Hsp104^K358D-Y257T^ (right). Nematodes have six anterior DA neurons (4 CEP and 2 ADE), which were scored at day 7 and 10 post-hatching. Left, the worm has only two normal neurons (2 CEP), where the other four neurons have degenerated. Right, the full complement of six anterior DA neurons expressing Hsp104^K358D-Y257T^ + α-syn, indicating a protective activity against α-syn toxicity. Triangles show normal neurons while arrows depict regions where there are degenerating or missing neurons. See also **Figure S6**.

Remarkably, α-syn-specific variant, Hsp104^K358D:Y257T^, strongly protected against α-syn-induced neurodegeneration at days 7 and 10 (Figure 7A, B). Despite only moderate α-syn toxicity suppression in yeast, Hsp104^K358D:Y257T^ displayed the strongest protection in the context of a metazoan nervous system, as ∼43% of worms retained the full complement of DA neurons at day 7, and ∼27% at day 10 (Figure 7A, B). Hsp104^K358D:Y257T^ is more effective in the *C. elegans* PD model than previously engineered potentiated variants Hsp104^A503S^ and Hsp104^DPLF:A503V^ (Figure 7A, B) (Jackrel et al., 2014a). Hsp104^K358D:Y257T^ expressed lower mRNA levels than generally potentiated variants in transgenic *C. elegans*, and did not affect α-syn expression levels (Figure S6). This finding suggested that the reduced neurodegeneration is due to the enhanced activity of this α-syn-specific Hsp104 variant.

Interestingly, another α-syn-specific variant, Hsp104^K358D:Y662M^, conferred neuroprotection but was only significantly protective against neurodegeneration at a later timepoint (day 10), where ∼30% of worms retained the full complement of WT neurons (Figure 7A). As Hsp104^K358D:Y662M^ did not prevent neurodegeneration at an earlier time point (day 7), this variant did not appear able to halt disease onset, but instead limited the progression of α-syn neurotoxicity at a threshold of greater severity. Delayed prevention of α-syn-induced neurodegeneration could stem from a unique remodeling activity of Hsp104^K358D:Y662M^. Using an Hsp104 variant that stops disease progression earlier (Hsp104^K358D:Y257T^), followed by Hsp104^K358D:Y662M^ which works more effectively later in disease, may be an advantageous combination strategy for mitigating α-syn toxicity in dopaminergic neurons.

## Discussion

Here, we engineered Hsp104 variants that selectively suppress toxicity of α-syn. Unexpectedly, the Hsp104 background used to engineer enhanced substrate specificity was critical. For example, we found that Hsp104^A503S^, a potentiated Hsp104 variant that breaks MD-MD contacts (Tariq et al., 2019; Ye et al., 2020), was not amenable for introducing substrate specificity via mutation of substrate-binding, pore-loop tyrosines. By contrast, α-syn-selective Hsp104 variants could be isolated by mutating pore-loop tyrosines in enhanced backgrounds that instead break NBD1-MD contacts, e.g. Hsp104^K358D^ or Hsp104^D484K^ (Lipinska et al., 2013). Thus, the precise nature of the underlying potentiation mechanism is an important aspect to consider when tuning Hsp104 variants (Ye et al., 2020).

By making subtle changes to the substrate-binding, pore-loop tyrosines that line the axial channel of Hsp104, we re-wired enhanced Hsp104 variants (Hsp104^K358D^ or Hsp104^D484K^) into a substrate-specific variants. Thus, we shifted the substrate repertoire of Hsp104 such that α-syn toxicity could be mitigated, but TDP-43 or FUS toxicity could not. α-Syn-selectivity could be conferred by Y257I/V/T/Q mutations in pore-loop 1 or by Y662L/M mutations in pore-loop 2 in the K358D background. Surprisingly, two classes of α-syn-specific Hsp104 variant emerged that reduced α-syn toxicity via distinct mechanisms.

The first class, which includes Hsp104^K358D:Y662M^ and Hsp104^K358D:Y257L:Y662M^, mitigated α-syn toxicity via ATPase-dependent disaggregation of α-syn inclusions. Similar to our previously engineered, potentiated Hsp104 variants, Hsp104^K358D:Y662M^ and Hsp104^K358D:Y257L:Y662M^ exhibited elevated ATPase activity compared to Hsp104. Hsp104^K358D:Y662M^ and Hsp104^K358D:Y257L:Y662M^ cleared α-syn foci in yeast, but did not affect TDP-43 or FUS aggregation. Importantly, Hsp104^K358D:Y662M^ dissolved α-syn fibrils but not TDP-43 fibrils *in vitro*. Thus, these Hsp104 variants have altered substrate specificity, which reduces disaggregase activity against some substrates (e.g. TDP-43) but permits effective α-syn disaggregation.

The second class, which includes Hsp104^K358D:Y257T^, mitigated α-syn toxicity via ATPase-dependent detoxification of α-syn conformers without disaggregation. Intriguingly, Hsp104^K358D:Y257T^ exhibited similar ATPase activity to WT Hsp104. However, Hsp104^K358D:Y257T^ was unable to disaggregate luciferase *in vitro*, but could confer some level of thermotolerance *in vivo* indicating that it retains some disaggregase activity. Unexpectedly, Hsp104^K358D:Y257T^ neither eliminated α-syn foci in yeast nor disaggregated α-syn fibrils *in vitro*. Rather, unlike WT Hsp104, Hsp104^K358D:Y257T^ colocalized with α-syn foci. We suggest Hsp104^K358D:Y257T^ may partially remodel α-syn inclusions into less toxic structures. Hsp104^K358D:Y257T^ could extract essential proteins trapped in α-syn foci or mitigate toxic interactions between α-syn and intracellular vesicles or organelles (Gitler et al., 2008; Mahul-Mellier et al., 2020; Shahmoradian et al., 2019; Soper et al., 2008).

Hsp104 variants from each class (Hsp104^K358D:Y662M^ and Hsp104^K358D:Y257T^) prevented dopaminergic neuron degeneration in a *C. elegans* model of PD more effectively than previous non-α-syn-specific, enhanced Hsp104 variants. Hsp104^K358D:Y257T^ protected against neurodegeneration throughout the course of disease in this model, whereas Hsp104^K358D:Y662M^ protected more strongly at later time points. Thus, substrate-specific Hsp104 variants effectively suppress neurodegeneration in the context of an intact metazoan nervous system, thereby translating the therapeutic benefits of these variants from yeast to metazoa. These findings bode well for subsequent validation, given the proven translational efficacy of the *C. elegans* α-syn model used in these studies, which has reproducibly demonstrated a predictive capacity to yield outcomes representative of mammalian models of PD (Gaeta et al., 2019).

We have also established that the level of off-target toxicity in yeast inherent to potentiated Hsp104 variants does not necessarily predict whether these variants will be neuroprotective in *C. elegans*. For example, Hsp104^K358D:Y662M^ exhibited off-target toxicity in yeast, but mitigated α-syn-induced dopaminergic neurodegeneration at later time points in *C. elegans*. Hsp104^K358D:Y662M^ could target essential yeast proteins for unfolding, but these proteins may not be as crucial or may be absent from *C. elegans*. By contrast, Hsp104^K358D:Y257L:Y662M^ exhibited off-target toxicity in yeast but did not prevent neurodegeneration in *C. elegans*. The challenge of translating our findings from yeast to worm likely reflects key differences between the two model systems. One important distinction is that the proteome of dopaminergic neurons in *C. elegans* likely has crucial differences from the yeast proteome. For example, dopamine modulates the propensity of α-syn to oligomerize and impacts neurodegeneration in mice, dopaminergic neuron culture, and *C. elegans* (Conway et al., 2001; Mor et al., 2017). Thus, the absence of dopamine in yeast, as well different sets of Hsp104-interacting proteins in yeast versus worm renders the direct translation of Hsp104 variants to metazoa challenging. Nevertheless, our efforts to engineer α-syn-specific Hsp104 variants have yielded Hsp104^K358D:Y662M^ and Hsp104^K358D:Y257T^, which outperform prior potentiated variants. It will be important to advance these Hsp104 variants to mammalian models of α-synucleinopathies.

It will also be of great interest to engineer substrate-specific Hsp104 variants to specifically target a range of other misfolded proteins in neurodegenerative disease, including TDP-43 and FUS. At a minimum, our findings suggest that engineering substrate-specific disaggregases can improve their ability to confer neuroprotection. We envision that increasing the substrate specificity of enhanced disaggregases could be applied broadly to tailor therapeutics for neurodegenerative disease. Beyond Hsp104, we anticipate that fine-tuning disaggregases found in humans, such as Hsp110, Hsp70, and Hsp40, nuclear-import receptors, or Skd3 will also be immensely valuable (Cupo and Shorter, 2020; Guo et al., 2019; Guo et al., 2018; Mack and Shorter, 2016; Shorter, 2011, 2017). We suggest that highly tuned, specific disaggregases that reverse targeted toxic misfolding events represent an exciting avenue for therapeutic agents in neurodegenerative disease (Mack and Shorter, 2016; Shorter, 2017).

## Acknowledgements

We thank Ben Deverett for assistance with analysis of microscopy. K.L.M. was supported by an NSF graduate research fellowship (DGE-1321851). M.E.J. was supported by NIH grant R35GM128772. J.L. was supported by an Alzheimer’s Association Research Fellowship. E.C. was supported by NIH grant T32GM008076 and a Blavatnik Family Fellowship in Biomedical Research. R.R.C. was supported by NIH grants T32GM008275, F31AG060672, and a Blavatnik Family Fellowship in Biomedical Research. L.M.C. was supported by an NSF graduate research fellowship (DGE-0822). J.S. was supported by NIH grants (DP2OD002177, R01GM099836), a Muscular Dystrophy Association Research Award (MDA277268), an ALS Association Award, the Life Extension Foundation, a Linda Montague Pechenik Research Award, the Packard Center for ALS Research at Johns Hopkins University, and Target ALS.

## Methods

### Yeast strains, media, and plasmids

Yeast strains used were wild-type W303a (*MATa, can1-100, his3-11, 15, leu2-3, 112, trp1-1, ura3-1, ade2-1*) or the isogenic strain W303aΔ*hsp104* (Jackrel et al., 2014a). The yeast strains W303aΔ*hsp104*-pAG303GAL-α-syn-YFP-pAG304GAL-α-syn-YFP, W303aΔ*hsp104*-pAG303GAL-FUS, W303aΔ*hsp104*-pAG303GAL-TDP-43, W303aΔ*hsp104*-pAG303GAL-TDP-43-GFPS11-pAG305GAL-GFPS1-10, and W303aΔ*hsp104*-pAG303GAL-FUS-GFP have been described previously (Jackrel et al., 2014a; Jackrel and Shorter, 2014a; Jackrel et al., 2014b). Yeast were grown in rich medium (YPD) or in synthetic media without amino acids used for selection. 2% sugar (dextrose, raffinose, or galactose) was added to synthetic media. Hsp104 variants were under control of a galactose-inducible promoter on pRS416GAL plasmids, except for in thermotolerance assays where they were under control of the HSE promoter on pRS416HSE plasmids. For mCherry-tagged Hsp104 variants, mCherry is located at the C-terminal end of Hsp104, separated by a glycine-serine linker (also on pRS416GAL plasmids).

### Generation of Hsp104 variants

Mutations were introduced into Hsp104 through QuikChange site-directed mutagenesis (Agilent) and confirmed by DNA sequencing.

### Yeast transformation and spotting assays

Plasmids containing Hsp104 variants were transformed into yeast using a standard lithium acetate and polyethylene glycol procedure (Gietz and Schiestl, 2007). For spotting assays, yeast cultures were grown to saturation overnight at 30°C in dropout media containing raffinose. Raffinose cultures were then normalized according to OD_600_, five-fold serial diluted, and spotted onto glucose and galactose plates using a 96-bolt replicator tool. Plates were grown at 30°C for 2-3 days and imaged.

### Toxicity spotting assay

pRS416GAL plasmids containing Hsp104 variants were transformed into W303aΔ*hsp104* yeast. Yeast cultures were grown to saturation overnight at 30°C in dropout media containing raffinose. Raffinose cultures were then normalized according to OD_600_ and five-fold serial diluted. The cultures were spotted onto two sets of glucose and galactose plates using a 96-bolt replicator tool. One set of plates was grown at 30°C, and the other at 37°C, for 2-3 days and subsequently imaged.

### Western blotting

Hsp104 variants transformed into appropriate yeast strains were grown to saturation overnight at 30°C in dropout media containing raffinose. Cultures were normalized to OD_600_ = 0.3 and grown in galactose dropout media at 30°C to induce Hsp104 and disease substrate expression (α-syn cultures induced for 8h, TDP-43 and FUS cultures induced for 5h). Galactose cultures were then normalized according to OD_600_ and the equivalent of 6ml culture with an OD_600_ = 0.6 were harvested by centrifugation. Media was aspirated, and the cell pellets were resuspended in 0.1M NaOH and incubated at room temperature for 5min. Cells were pelleted again by centrifugation, supernatant removed, and pellet was resuspended in 100μL 1X SDS sample buffer and boiled for 4-5min. Samples were separated via SDS-PAGE (4-20% gradient, Bio-Rad) and transferred to a PVDF membrane (Millipore) using a Trans-Blot SD Semi-Dry Transfer Cell (Bio-Rad). Membranes were blocked for at least 1h at room temperature and then incubated with primary antibodies (rabbit anti-Hsp104 polyclonal (Enzo Life Sciences); rabbit anti-FUS polyclonal (Bethyl Laboratories); rabbit anti-TDP-43 polyclonal (Proteintech); rabbit anti-GFP polyclonal (Sigma-Aldrich); mouse anti-PGK1 monoclonal (Invitrogen)) at 4°C overnight. Membranes were washed multiple times with PBS-T, incubated with secondary antibodies (goat anti-mouse and goat anti-rabbit, LI-COR) for 1h at room temperature, and washed again multiple times with PBS-T (final wash with PBS). Membranes were imaged using a LI-COR Odyssey FC Imaging system.

### Thermotolerance assay

Hsp104 variants under the HSE promoter were transformed into W303aΔ*hsp104* yeast. Yeast cultures were grown to saturation overnight at 30°C in glucose dropout media. Cultures were normalized to OD_600_ = 0.3 and grown in glucose dropout media at 30°C for at least 4h, after which the equivalent of 6 ml culture with an OD_600_ = 0.6 was grown at 37°C for 30 min (if assessing Hsp104 expression, samples would be harvested at this stage for western blot as described above). Cultures were then heat-shocked at 50°C in 1.5ml Eppendorf tubes in an Eppendorf Thermomixer for 30min and incubated on ice for 2min. Cultures were diluted appropriately, plated on glucose dropout media, and incubated at 30°C. After 2-3 days, colonies were counted using an aCOLyte colony counter and software (Synbiosis). W303a yeast were transformed with an empty pRS416Gal vector and treated as above for endogenous Hsp104 control.

### Fluorescence microscopy

Yeast strains used in fluorescence microscopy in (Jackrel et al., 2014a) were used. Hsp104 variants on pRS416GAL plasmids were transformed into each yeast strain (α-syn-YFP, FUS-GFP, or TDP43-GFPS11 from (Jackrel et al., 2014a)) and grown to saturation overnight at 30°C in dropout media containing raffinose. Cultures were normalized to OD_600_ = 0.3 and grown at 30°C in galactose dropout media to induce Hsp104 and disease substrate expression (α-syn cultures induced for 8h, TDP-43 and FUS cultures induced for 5h). For α-syn and FUS cultures, cells were harvested and processed for microscopy, and imaged as live cells. For TDP-43 cultures, cells were harvested and fixed by spinning cells down, resuspending in 1ml cold 70% ethanol, and immediately pelleting cells again. Cells were washed 3 times with cold PBS and resuspended in Vectashield mounting medium with 4’,6-diamidino-2-phenylindole (DAPI) (Vector Laboratories). Cells were imaged at 100X magnification using a Leica DM IRBE microscope. Cells were analyzed using ImageJ software, and a minimum of 200 cells were quantified per sample in at least three independent trials. One-way ANOVA with Dunnett’s *post hoc* test was performed using GraphPad Prism Software.

For fluorescence microscopy to assess mCherry-GFP localization, Hsp104 variants with a C-terminal mCherry tag, or mCherry alone (on pRS416GAL plasmids) were transformed into the α-syn-YFP strain described in (Jackrel et al., 2014a). Cultures were treated as α-syn cultures described above. After induction, cells were harvested, stained with Hoechst dye (channel not shown in Figure 5C) and imaged at 100X magnification using a Leica DM IRBE microscope. For analysis of mCherry colocalization with GFP, cells were imaged in red (TX2) and green (L5) channels. A semi-automated procedure was used to quantify colocalization between the red and green images (code available at https://github.com/bensondaled/mack-cell-annotation). Cells containing α-syn-YFP foci were manually identified and selected using a custom interface. A colocalization metric for each cell was then defined as the Pearson correlation between the pixels in the red and green channels. A cell was defined to exhibit colocalization if this measure exceeded 0.85, a uniform threshold applied to all images and chosen for its agreement with manual colocalization assessments. A minimum of 200 cells were quantified per sample in at least three independent trials. One-way ANOVA with Dunnett’s *post hoc* test was performed using GraphPad Prism Software.

### *C. elegans* dopaminergic neurodegeneration

Nematodes were grown and maintained using well-established procedures (Brenner, 1974). Plasmid constructs were injected into worms to generate transgenic animals using previously described methods (Berkowitz et al., 2008). Strains UA367 (*baEx198 a,b,c* [P_*dat-1*_::HSP104-K358D, *rol-6*];*baln11* [P_*dat-1*_ :: α-syn, P_*dat-1*_ ::GFP]), UA368 (*baEx199 a,b,c* [P_*dat-1*_::HSP104-K358D:Y257L, *rol-6*];*baln11* [P_*dat-1*_ :: α-syn, P_*dat-1*_ ::GFP]), UA369 (*baEx200 a,b,c* [P_*dat-1*_::HSP104-K358D:Y257T, *rol-6*];*baln11* [P_*dat-1*_ :: α-syn, P_*dat-1*_ ::GFP]), UA370 (*baEx201 a,b,c* [P_*dat-1*_::HSP104-K358D:Y257L:Y662M, *rol-6*];*baln11* [P_*dat-1*_ :: α-syn, P_*dat-1*_ ::GFP]), and UA371 (*baEx202 a,b,c* [P_*dat-1*_::HSP104-K358D: Y662M, *rol-6*];*baln11* [P_*dat-1*_ :: α-syn, P_*dat-1*_ ::GFP]) were generated by injecting 50 ng/µl of corresponding plasmid construct into UA44 (*baln11* [P_*dat-1*_ :: α-syn, P_*dat-1*_ ::GFP]) with phenotypic marker (*rol-6*, 50 ng/µl, for roller expression). Three independent stable lines were created for each group (*a, b, c*). Strain UA256 (*baEx147 a,b,c* [P_*dat-1*_::Hsp104 WT, P_*myo-2*_::mCherry]; *baIn11* [P_*dat-1*_ :: α-syn, P_*dat-1*_ ::GFP]), UA259 (*baEx150 a,b,c* [P_*dat-1*_::Hsp104-A503S, P_*myo-2*_::mCherry]; *baIn11* [Pdat-1:: α-syn, P_*dat-1*_::GFP]), and UA262 (*baEx153 a,b,c* [P_*dat-1*_::Hsp104-DPLF:A503V, P_*myo-2*_::mCherry]; *baIn11*[P_*dat-1*_:: α-syn, P_*dat-1*_::GFP]) were previously generated (Jackrel et al., 2014a).

For dopaminergic neurodegeneration analyses, the transgenic animals were scored as described previously (Hamamichi et al., 2008). Briefly, on the day of analysis, the six anterior dopaminergic neurons [four CEP (cephalic) and two ADE (anterior deirid)] were examined in 30 animals randomly. Each nematode was considered normal when all six anterior DA neurons were present. However, if a worm exhibited any degenerative phenotype, such as a missing dendritic process, cell body loss, or a blebbing neuronal process, it was scored as degenerating. Three independent transgenic worm lines were analyzed per genetic background for a total of ninety animals per transgenic line and an average of total percentage of worms with normal neurons was reported in the study. A one-way ANOVA, followed by a Dunnett’s *post hoc* test, was performed for statistical analysis using GraphPad Prism Software.

### Quantitative Real-Time PCR in nematodes

RNA isolation and qRT-PCR were performed on worms using previously published methods (Hamamichi et al., 2008; Kim et al., 2018). Briefly, total RNA was isolated from 100 young adult (day 4 post hatching) nematodes from corresponding transgenic line using TRI reagent (Molecular Research Center). 1µg of RNA was used for cDNA synthesis using the iScript Reverse Transcription Supermix for qRT-PCR (Bio-Rad, Hercules, CA, USA). qRT-PCR was performed using IQ-SYBR Green Supermix (Bio-Rad) with the Bio-Rad CFX96 Real-Time System. Hsp104 expression levels were normalized to two reference genes (*snb-1* and *y45*). α-syn expression levels were normalized to three reference genes (*ama-1, cdc-42*, and *tba-1*). The reference target stability was analyzed by GeNorm and passed for all reference genes listed above. Three technical replicates with three biological replicates were performed in this study. Each primer pair was confirmed for at least 90-100% efficiency by standard curve analyses. The following primers were used for the assays:

*hsp104* Forward: GGCCATGAAGAATTGACACAAG

*hsp104* Reverse: CCTTAAATCAGCGGCGGTAG

*α-syn* Forward: ATGTAGGCTCCAAAACCAAGG

*α-syn* Reverse: ACTGCTCCTCCAACATTTGTC

*snb-1* Forward: CCGGATAAGACCATCTTGACG

*snb-1* Reverse: GACGACTTCATCAACCTGAGC

*y45* Forward: GTCGCTTCAAATCAGTTCAGC

*y45* Reverse: GTTCTTGTCAAGTGATCCGACA

*ama-1* Forward: TCCTACGATGTATCGAGGCAA

*ama-1* Reverse: CTCCCTCCGGTGTAATAATGA

*cdc-42* Forward: CCGAGAAAAATGGGTGCCTG

*cdc-42* Reverse: TTCTCGAGCATTCCTGGATCAT

*tba-1* Forward: ATCTCTGCTGACAAGGCTTAC

*tba-1* Reverse: GTACAAGAGGCAAACAGCCAT

### Protein purification

#### Hsp104 and HAP

Hsp104 and HAP proteins were purified as previously described (Jackrel et al., 2014a) with the following modifications. Eluate from Affi-Gel Blue Gel was equilibrated to a low-salt buffer Q (∼100mM NaCl, 20mM TRIS pH 8.0, 5mM MgCl_2_, 0.5mM EDTA) and purified via ResourceQ anion exchange chromatography. Buffer Q (20mM TRIS pH 8.0, 50mM NaCl, 5mM MgCl_2_, 0.5mM EDTA) was used as running buffer, and the protein was eluted with a linear gradient of buffer Q+ (20mM TRIS pH 8.0, 1M NaCl, 5mM MgCl_2_, 0.5mM EDTA). The eluted protein was buffer-exchanged into high-salt storage buffer (40mM HEPES-KOH pH 7.4, 500mM KCl, 20mM MgCl_2_) plus 10% glycerol and 1mM DTT and snap-frozen.

#### GroEL_trap_

pTrc99A-GroEL_trap_ was transformed into DH5α competent *E. coli* cells (Thermo Fisher). Cells were grown in 2xYT medium with appropriate antibiotics at 37°C with shaking until OD_600_ reached ∼ 0.4 – 0.6. Protein overexpression was induced with 1mM IPTG, and cells were grown at 37°C until OD_600_ ∼ 2.0. Cells were harvested by spinning (5000g, 4°C, 15 min) and pellet was resuspended in 50mM sodium phosphate buffer and centrifuged (5000g, 4°C, 15min). Pellet was resuspended in low-salt buffer (50mM Tris-HCl pH 7.5, 1mM EDTA, 1mM DTT, 50mM NaCl) and 10mg lysozyme per g cell pellet. Sample stirred gently for 5min, lysed through sonication, and centrifuged (17,000 g, 4°C, 30min). Clarified lysate loaded onto HiTrap Q HP column (GE Healthcare) and eluted through salt gradient using low-salt buffer (as described above) and high-salt buffer (50mM Tris-HCl pH 7.5, 1mM EDTA, 1mM DTT, 500mM NaCl (Braig et al., 1994). Collected fractions were exchanged into the following TKME-100 buffer: 20mM TRIS-HCl pH 7.5, 100mM KCl, 10mM MgCl_2_, 0.1mM EDTA, 5mM DTT, 10% glycerol, 0.005% Triton X-100 and snap-frozen.

#### RepA_1-25_-GFP

pBAD-RepA_1-25_-GFP was transformed into BL21 (DE3)-RIL cells. Cells were inoculated in 2xYT medium with appropriate antibiotics at 37°C with shaking until OD_600_ reached ∼ 0.6 – 0.8. Protein overexpression was induced with 1mM IPTG, and cells were grown at 30°C for 4h. Cells were harvested by spinning (4000rpm, 25min) and pellet was resuspended in 40mM HEPES-KOH pH 7.4 plus 2mM 2-Mercaptoethanol (BME) and EDTA-free protease inhibitors. Cells were lysed using a sonicator and centrifuged (16,000rpm, 20min). The resulting pellet was washed twice with HM buffer (40mM HEPES-KOH pH 7.4, 20mM MgCl_2_) plus 2mM BME. After each wash, cells were centrifuged (16,000 rpm, 20min). Pellet was then resuspended in buffer containing 8M urea, 40mM Tris-HCl pH 6.8, 500mM NaCl, 10% glycerol (v/v) and agitated slowly overnight at 25°C. Sample was centrifuged (16,000rpm, 20min) and supernatant was kept. Supernatant was incubated with Ni-NTA beads (HisPur™ Ni-NTA Resin, Thermo Scientific) pre-equilibrated in buffer containing 8M urea, 40mM Tris pH 6.8, 500mM NaCl, 10% glycerol (v/v) for 2h on a spinning wheel at 25°C at the lowest speed. Sample washed 5 times with buffer containing 8M urea, 40mM Tris-HCl pH 6.8, 500mM NaCl, 20mM imidazole, 10% glycerol (v/v). Sample then washed 5 times with buffer containing 8M urea, 40mM Tris-HCl pH 6.8, 500mM NaCl, 40mM imidazole, 10% glycerol (v/v). Sample eluted with buffer containing 8 M urea, 40mM Tris-HCl pH 6.8, 500mM NaCl, 500mM imidazole, 10% glycerol (v/v). Sample was dialyzed overnight into buffer containing 40mM HEPES-KOH pH 7.4, 20mM imidazole, 150mM KCl, 2mM BME, 10% glycerol (v/v) at 4°C and re-loaded onto a Ni-NTA column (HisTrap™ HP, GE Healthcare). An imidazole gradient was applied (from 20mM to 500mM) over 20CV in buffer containing 40mM HEPES-KOH pH 7.4, 20mM imidazole, 150mM KCl, 2mM BME, 10% glycerol (v/v). The purity of eluted fractions was assessed using SDS-PAGE. Collected fractions were buffer-exchanged into HKM-150 buffer (40mM HEPES-KOH pH 7.4, 150mM KCl, 20mM MgCl_2_) plus 2 mM BME and 10% glycerol (v/v) and snap-frozen.

#### ClpP

C-terminally His-tagged ClpP was purified as previously described (Jackrel et al., 2014a).

#### Hsp70 and Hsp40

Hsc70 and Hdj2 were from Enzo Life Sciences. Hsp72 and Hdj1 were purified as in (Michalska et al., 2019).

#### α-Syn and TDP-43

α-Syn and TDP-43 were purified as described (Jackrel et al., 2014a).

#### ATPase assay

Hsp104 (0.25μM monomer in Hsp104 buffer, see Hsp104 purification) was incubated with ATP (1 mM) for 5min at 25°C in luciferase-refolding buffer (LRB: 25mM HEPES-KOH pH 7.4, 150mM KAOc, 10mM MgAOc, 10mM DTT). The final reaction buffer contained 6.3% of HKM-500 buffer. ATPase activity was evaluated by the release of inorganic phosphate, which was measured using a malachite green phosphate detection kit (Innova Biosciences). Background hydrolysis at time zero was subtracted.

### FITC-casein degradation assay

FITC-casein (0.1μM) was incubated with HAP or HAP variants (1μM monomer), ClpP (21μM monomer), ATP (5mM), and an ATP regeneration system (1mM creatine phosphate and 0.25μM creatine kinase). ClpP was first buffer-exchanged into luciferase-refolding buffer (LRB: 25mM HEPES-KOH pH 7.4, 150mM KAOc, 10mM MgAOc, 10mM DTT) at 25°C. Reaction mixtures were assembled 25°C in LRB and degradation of FITC-casein was monitored by measuring fluorescence (excitation 490nm, emission 520nm) using a Tecan Infinite M1000 plate reader.

### RepA_1-25_-GFP unfoldase assay

RepA_1-25_-GFP (0.7μM) was incubated with Hsp104 or Hsp104 variants (2.1μM hexamer), ATP (4mM), ARS (20mM creatine phosphate, 0.06μg/μl creatine kinase). GroEL_trap_ (2.5μM tetradecamer) was included to prevent refolding of unfolded RepA_1-25_-GFP. Hsp104 variants were buffer-exchanged into TKME-100 buffer at 25°C. Reactions were assembled on ice in TKME-100 buffer plus 20μg/ml BSA. RepA_1-25_-GFP unfolding was measured by fluorescence (excitation 395nm, emission 510nm) using a Tecan Safire^2^, which was heated to 30°C prior to reading.

### Luciferase reactivation assay

Aggregated luciferase (100nM) was incubated with Hsp104 or Hsp104 variants (1μM monomer), ATP (5mM), and an ATP regeneration system (10mM creatine phosphate, 0.25μM creatine kinase) in the presence or absence of co-chaperones Hsp70 (Hsc70 or Hsp72, 0.167μM) and Hsp40 (Hdj2 or Hdj1, 0.167μM) for 90min at 25°C in LRB. The final reaction buffer contained 8.6% of HKM-500 buffer. After 90min, luciferase activity was measured with a luciferase assay reagent (Promega). Recovered luminescence was measured using a Tecan Infinite M1000 plate reader.

### α-Syn and TDP-43 fibril disaggregation

α-Syn (80μM) was assembled into fibrils via incubation in 40mM HEPES-KOH (pH 7.4), 150 mM KCl, 20 mM MgCl_2_, and 1 mM dithiothreitol for 48h at 37°C with agitation. α-Syn fibrils (3μM monomer) were incubated without or with Hsp104, Hsp104^A503S^, Hsp104^K358D^, Hsp104^K358D:Y257T^ or Hsp104^K358D:Y662M^ (3μM) plus Hsc70 (3μM), Hdj1 (3μM), ATP (20mM) and an ATP regeneration system (20mM creatine phosphate and 0.5μM creatine kinase) for 2h at 30°C. Disaggregation was assessed by Thioflavin-T fluorescence (Lo Bianco et al., 2008). To generate TDP-43 fibrils, GST-TEV-TDP-43 (6μM) was incubated with TEV protease in 40mM HEPES-KOH (pH 7.4), 150mM KCl, 20mM MgCl_2_, and 1mM dithiothreitol for 16h at 25°C with agitation, by which time all the TDP-43 had aggregated. TDP-43 fibrils (3μM monomer) were incubated without or with Hsp104, Hsp104^A503S^, Hsp104^K358D^, Hsp104^K358D:Y257T^ or Hsp104^K358D:Y662M^ (3μM) plus Hsc70 (3μM), Hdj1 (3μM), ATP (20 mM) and an ATP regeneration system (20mM creatine phosphate and 0.5μM creatine kinase) for 2h at 30°C. Disaggregation was assessed by turbidity (absorbance at 350nm).

---

**Figure S1.**
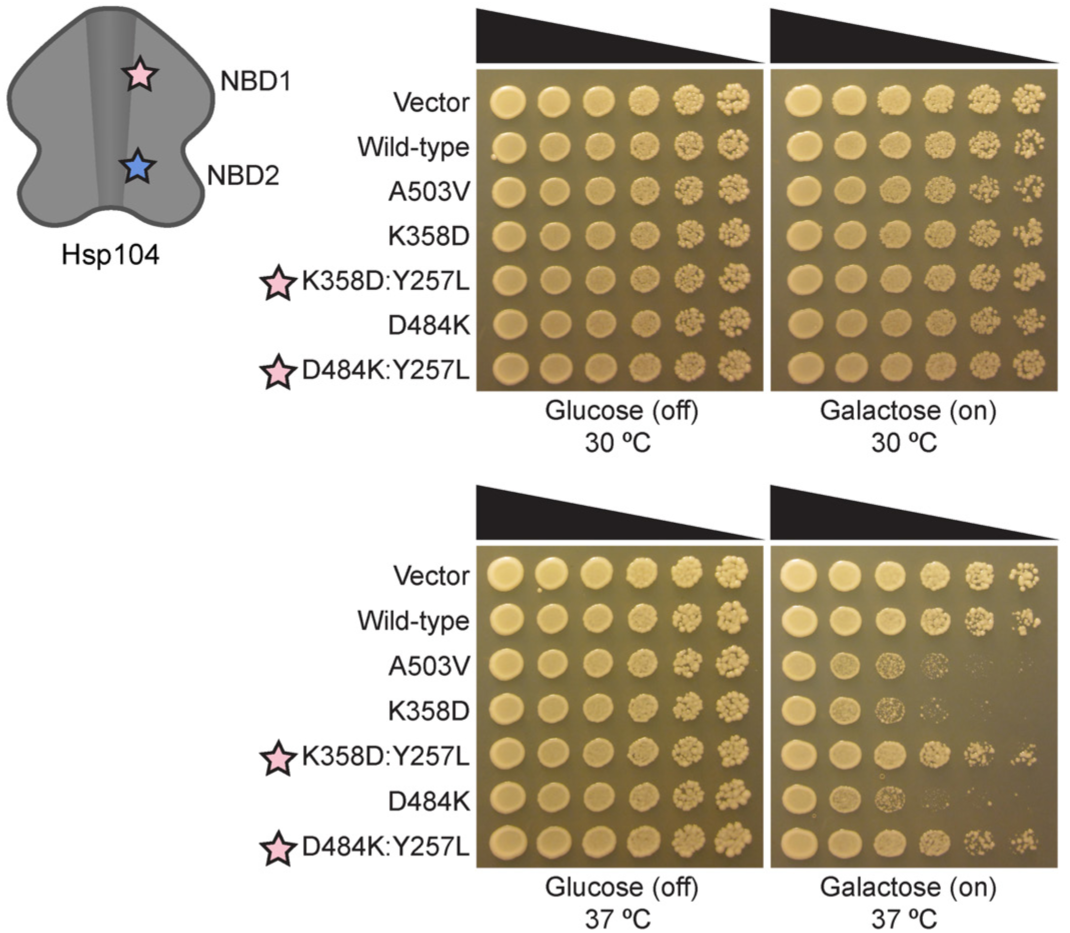
Hsp104^K358D^ and Hsp104^D484K^ are toxic to yeast at 37°C. Δ*hsp104* yeast were transformed with galactose-inducible Hsp104 variants or an empty vector and spotted onto glucose (uninducing, off) and galactose (inducing, on) media in a five-fold serial dilution. Yeast were incubated at 30°C (top) or 37°C (bottom). Stars indicate substitution to pore loop in NBD1 (pink), NBD2 (purple).

**Figure S2.**
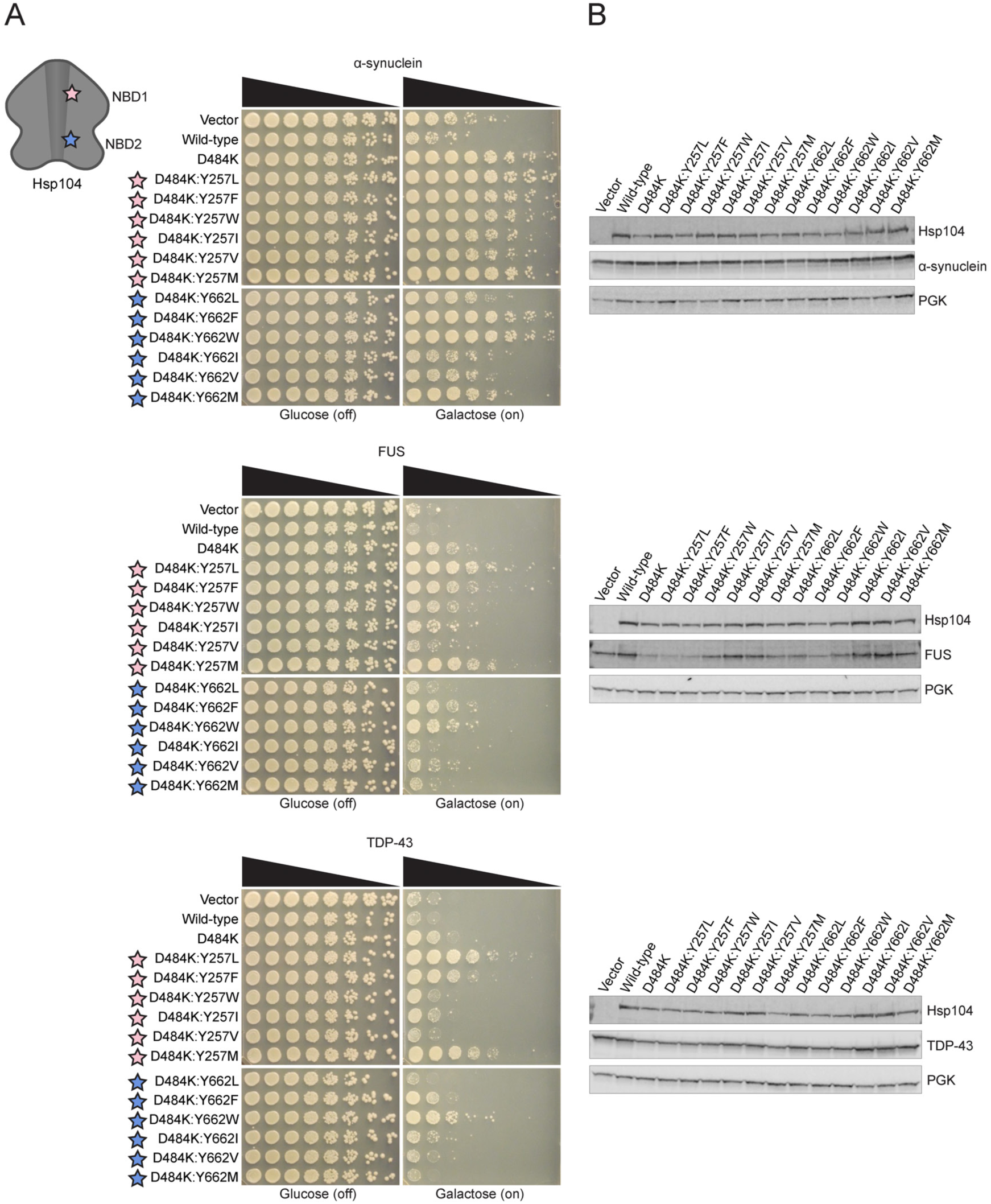
Hsp104^D484K^ pore loop variants have similar toxicity suppression profile to Hsp104^K358D^ variants. **(A**) Δ*hsp104* yeast integrated with α-syn-YFP (top), FUS (middle), or TDP-43 (bottom) on a galactose-inducible promoter were transformed with Hsp104 variants or an empty vector control. Yeast were spotted onto glucose (uninducing, off) and galactose (inducing, on) media in a five-fold serial dilution. Stars indicate substitution to pore loop in NBD1 (pink), NBD2 (purple). **(B)** Integrated strains from (A) were induced in the presence of Hsp104 variants or empty vector control for 5 hours (FUS, TDP-43) or 8 hours (α-syn). Yeast were lysed and lysates visualized via Western blot. 3-Phosphoglycerate kinase (PGK) is a loading control.

**Figure S3.**
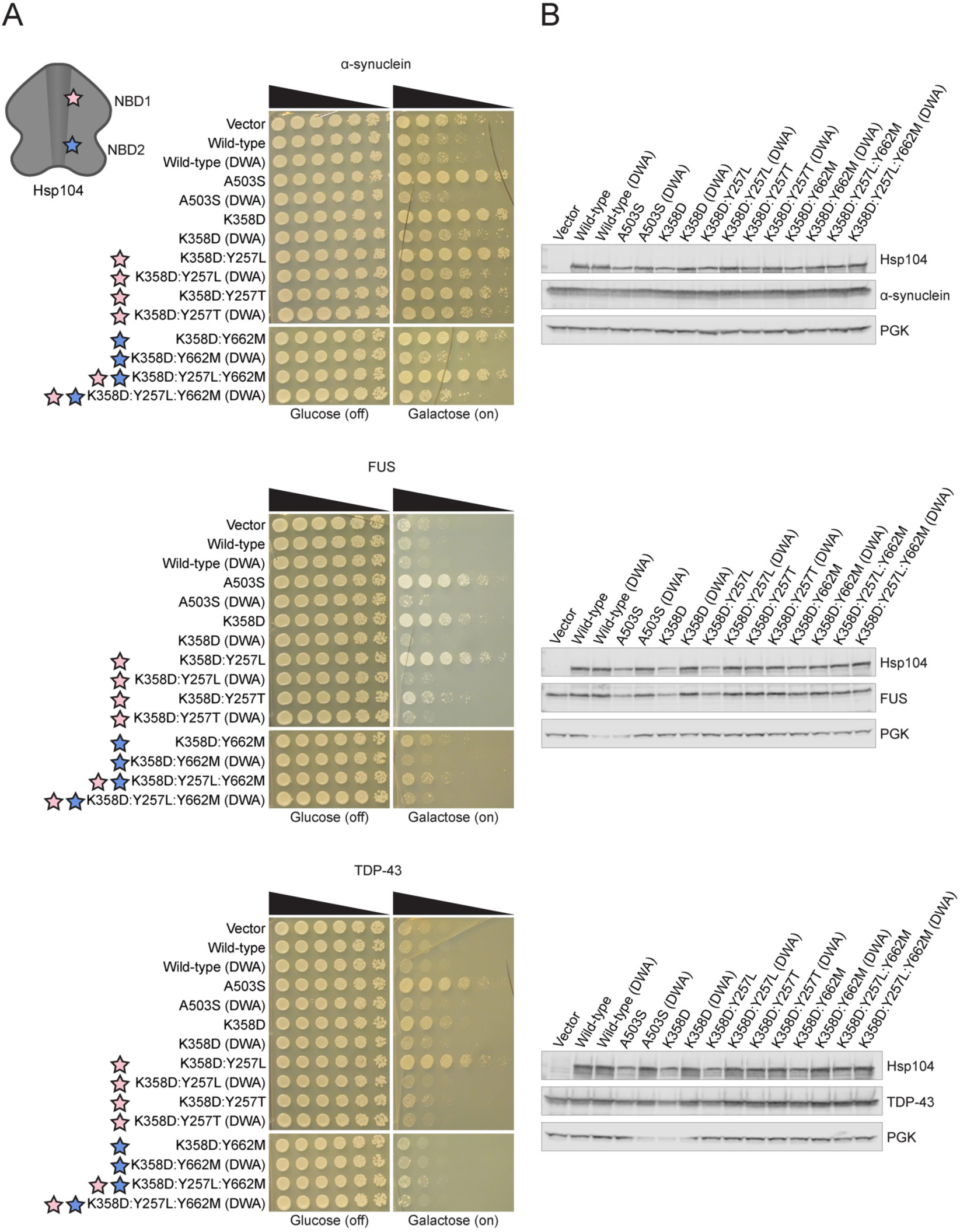
Hsp104^DWA^ pore-loop variants have diminished substrate toxicity suppression. **(A)** Δ*hsp104* yeast integrated with α-syn-YFP (top), FUS (middle), or TDP-43 (bottom) on a galactose-inducible promoter were transformed with Hsp104 variants or an empty vector control. Yeast were spotted onto glucose (uninducing, off) and galactose (inducing, on) media in a five-fold serial dilution. Stars indicate substitution to pore loop in NBD1 (pink), NBD2 (purple). **(B)** Integrated strains from (A) were induced in the presence of Hsp104 variants or empty vector control for 5 hours (FUS, TDP-43) or 8 hours (α-syn). Yeast were lysed and lysates visualized via Western blot. 3-Phosphoglycerate kinase (PGK) is a loading control.

**Figure S4.**
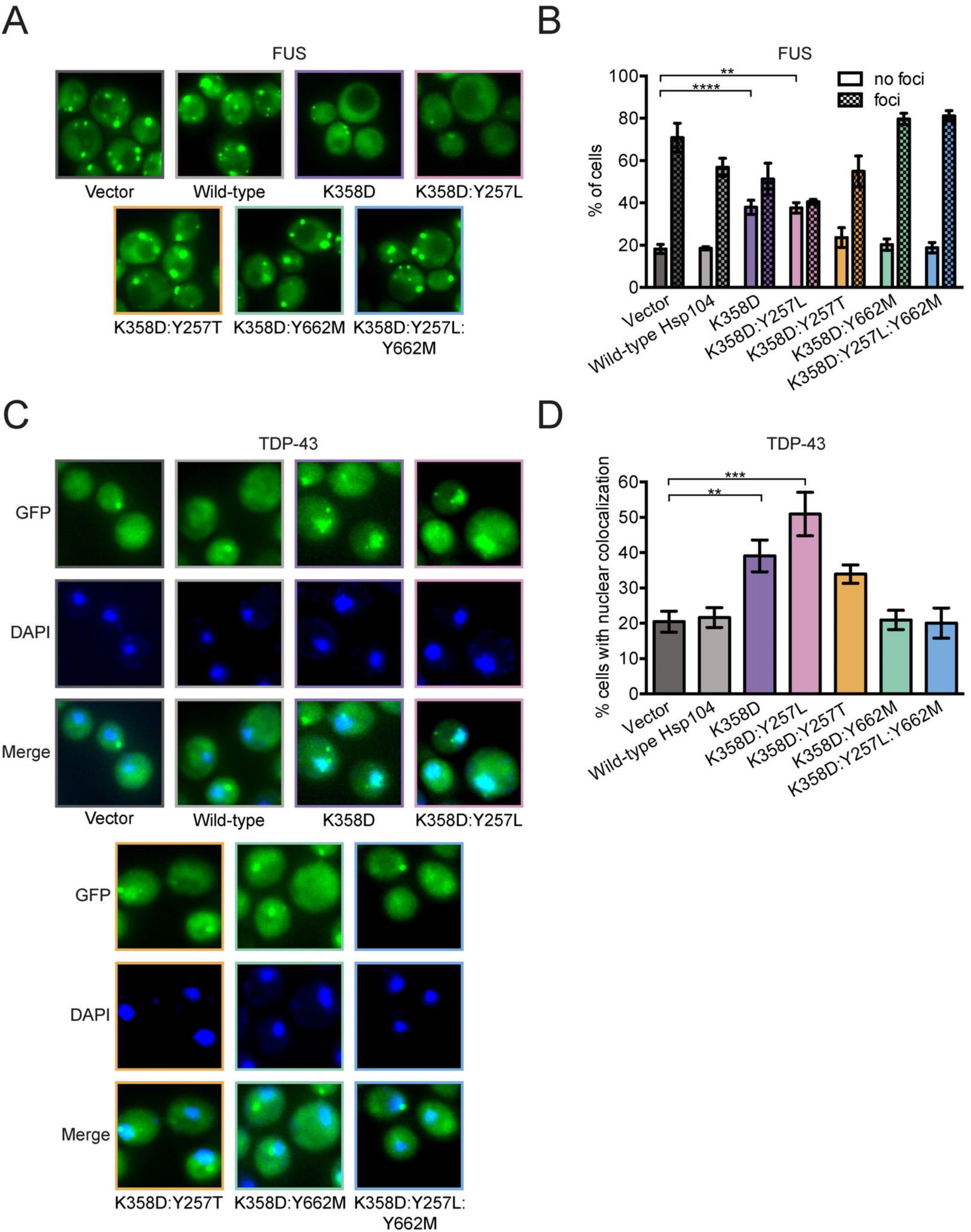
α-syn-specific Hsp104 variants do not significantly act on FUS or TDP-43 aggregates. **(A)** Representative fluorescence microscopy images of Δ*hsp104* yeast integrated with FUS-GFP and transformed with Hsp104 variants or an empty vector control. **(B)** FUS-GFP aggregation in yeast was quantified by counting the number of cells with FUS-GFP foci or no FUS-GFP foci. Values represent means ± SEM (n = 3-6). \*\**P*<0.01, \*\*\*\**P*<0.0001; One-way ANOVA with Dunnett’s *post-hoc* test. **(C)** Representative fluorescence microscopy images of Δ*hsp104* yeast integrated with TDP-43-GFPS11 and transformed with Hsp104 variants or an empty vector control. **(D)** TDP-43-GFPS11 localization in yeast was quantified by counting the number of cells with nuclear-localized TDP-43-GFPS11. Values represent means ± SEM (n = 3-6). \*\**P*<0.01, \*\*\**P*<0.001; One-way ANOVA with Dunnett’s *post-hoc* test.

**Figure S5.**
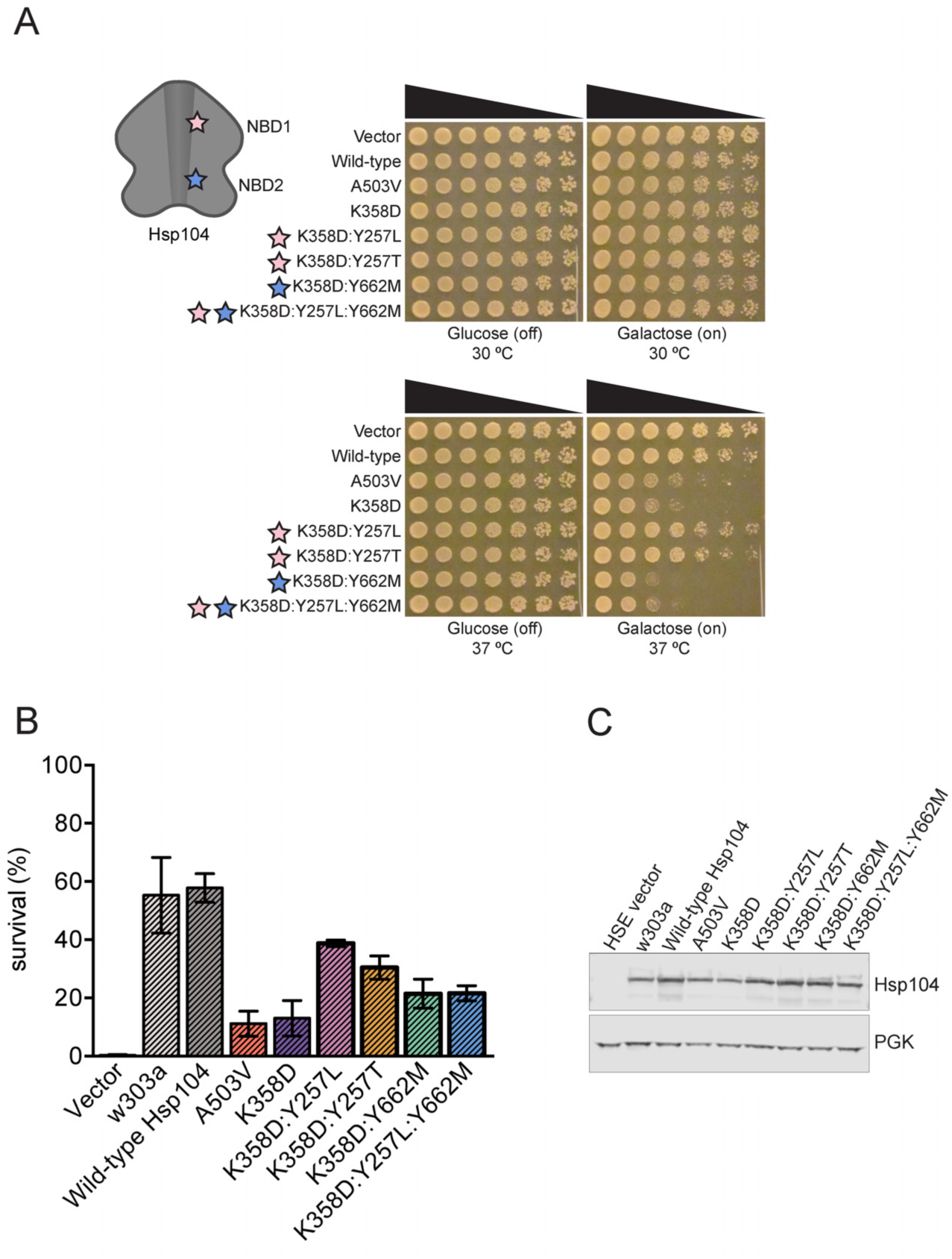
α-Syn-specific Hsp104 variants have inherently different toxicity in yeast and confer thermotolerance to different extents. **(A)** Δ*hsp104* yeast were transformed with galactose-inducible Hsp104 variants or an empty vector and spotted onto glucose (uninducing, off) and galactose (inducing, on) media in a five-fold serial dilution. Yeast were incubated at 30°C (top) or 37°C (bottom). Stars indicate substitution to pore loop in NBD1 (pink), NBD2 (purple). **(B)** Δ*hsp104* yeast were transformed with Hsp104 variants or an empty vector under the HSE promoter. Yeast were grown in selective media for 4 hours, and cultures were normalized and incubated at 37°C for 30 minutes. Cultures were then heat-shocked for 0, 20, or 30 minutes and plated on selective media. Plates were incubated at 30°C for ∼ 2 days and colonies counted. Wild-type w303a yeast expressing endogenous Hsp104 was included as a control. Values represent means ± SEM (n=3). **(C)** Yeast strains from (B) were grown in selective media for 4 hours. Cultures were normalized and grown for 30 minutes at 37°C to induce Hsp104 variant expression. Yeast were lysed, and lysates visualized via Western blot. 3-Phosphoglycerate kinase (PGK) is a loading control.

**Figure S6.**
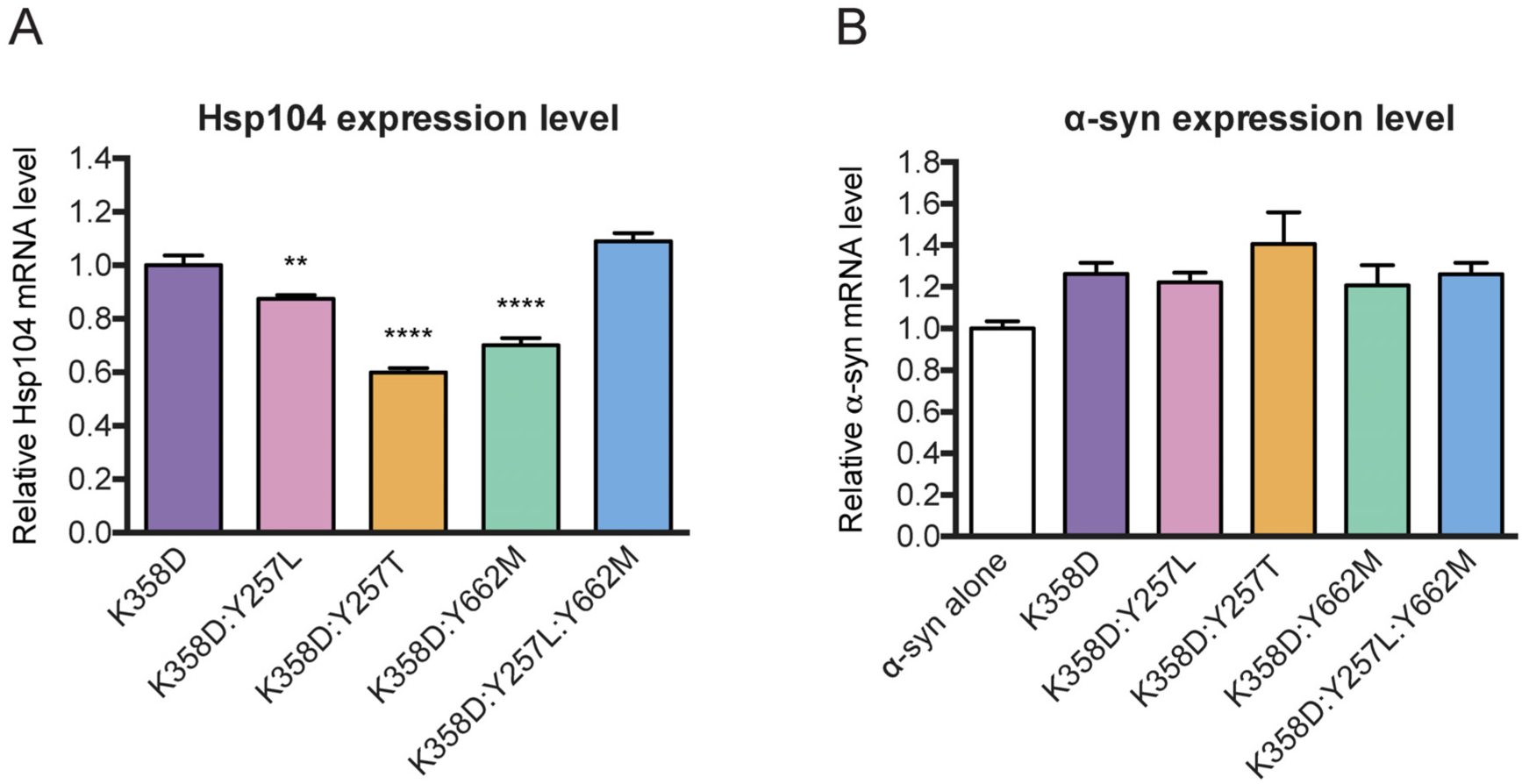
Confirmation of transcription levels of HSP104 variants and α-syn by qRT-PCR. **(A and B)** Transgenic *C. elegans* co-overexpressing Hsp104 variants + α-syn within DA neurons were subjected to qRT-PCR. **(A)** Hsp104 transgenic lines exhibit variable transcription levels. In this graph, all transgenic lines are displayed with their HSP104 mRNA levels relative to Hsp104^K358D^. Although Hsp104 variants Hsp104^K258D:Y257T^ and Hsp104^K358D:Y662M^ have lower expression levels than Hsp104^K358D^, they show greater protective activity against α-syn toxicity in Figure 7. **(B)** In this graph, all transgenic lines are displayed with their α-syn mRNA levels relative to *C. elegans* expressing α-syn without Hsp104. The overexpression of Hsp104 variants does not affect the transcription of α-syn, indicating that the observed neuroprotection is not caused by reduced α-syn expression. Values represent means ± SEM. \*\**P*<0.01, \*\*\*\**P*<0.0001; One-way ANOVA with Dunnett’s *post-hoc* test.

